# Radical fringe facilitates NOTCH1 and JAG1 *cis* interactions to sustain Hematopoietic stem cell fate

**DOI:** 10.1101/2023.04.19.537430

**Authors:** Roshana Thambyrajah, Maria Maqueda, Wen Hao Neo, Kathleen Imbach, Yolanda Guillen, Daniela Grases, Zaki Fadlullah, Stefano Gambera, Francesca Matteini, Xiaonan Wang, Fernando J. Calero-Nieto, Manel Esteller, Maria Carolina Florian, Eduard Porta, Rui Benedito, Berthold Göttgens, Georges Lacaud, Lluis Espinosa, Anna Bigas

## Abstract

Hematopoietic stem cells (HSCs) develop within a short time window from the hemogenic endothelium in the aorta- gonads-and mesonephros (AGM) region during embryonic development. The first HSCs reside within Intra-aortic hematopoietic clusters (IAHC) along with hematopoietic progenitors (HPC). The signalling mechanisms that divert HSCs from HPCs are unknown. Notch signaling is essential for arterial specification, IAHC formation and HSC activity, but current studies on how Notch drives these different fates are inconsistent. To determine the role of Notch in the specification of hemogenic endothelium, HSC and/or HPCs, we extensively analyzed Notch dynamics in the period of HSC generation. We defined the expression pattern of Notch signalling molecules at the gene and protein level and established a molecular mechanism that reconcile previous studies demonstrating the loss of HSC activity in NOTCH1, JAG1 and RBPJ null mutants, the enhanced HSC generation by blocking specific Notch activities or the abrogation of emerging HSCs by high Notch activation. We now demonstrate that Notch activity is highest in a subset of Gfi1+ hemogenic endothelial cells and is gradually lost with HSC maturation. We uncover that the HSC phenotype is maintained through loss of Notch activity due to increasing levels of NOTCH1 and JAG1 interactions on the surface of the same cell (*cis*) that renders the NOTCH1 receptor from being activated. Forcing activation of the NOTCH1 receptor in IAHC cells activates a hematopoietic differentiation program and supports a *cis*-inhibitory function for JAG1 and NOTCH1. Furthermore, we demonstrate that this *cis*-inhibitory interaction is enabled by RADICAL FRINGE (RFNG), a glycosyltransferase that enhances the affinity of NOTCH1 to JAG1 in *cis*. Finally, our results indicate that NOTCH1-JAG1 *cis*-inhibition is necessary for preserving the HSC phenotype in the hematopoietic clusters of the aorta.

## Introduction

Hematopoietic stem cells (HSCs) have the exceptional ability to self-renew and re-establish the entire blood system after injury or transplantation, making them very attractive to treat blood disorders. The initial HSCs are detected during mid-gestation (E10-E12) in the trunk of the embryo, where the aorta, gonads and mesonephros (AGM) co-localize. At the site of their origin, they are detected in intra-aortic hematopoietic clusters (IAHC), clusters of hematopoietic cells that reach into the lumen of the dorsal aorta (DA) ^1–4^. Although several hundreds of such hematopoietic cells (HSPC) are organized in 30-40 IAHC, only a small minority of these cells show HSC activity in transplantation assays ^5–7^. Markers that distinguish these early HSCs from multipotent progenitors (hereafter referred to as HPC) are only starting to emerge ^8^. Indeed, very recent studies demonstrate that the hematopoietic system with its established hierarchy, i.e., with long/short term (LT/ST-) HSCs and HPCs are already present in the AGM ^8–10^. The new findings suggests that all these hematopoietic cells migrate to the fetal liver to multiply simultaneously, as opposed to the (LT)-HSCs pool derived from the AGM generating the entire blood system in the fetal liver ^9, 10^. This new model implies that certain molecular mechanisms must be in place to support the emergence of these diverse blood populations within a short length of time in the AGM.

All IAHC are derived from hemogenic endothelial (HE) cells, a specialised endothelium that is embedded with the DA, but already expresses essential transcription factors of HSPC, including *Gata2*, *Runx1* and *Gfi1* ^11–16^. These HE cells undergo a transformation to a hematopoietic phenotype, termed endothelial to hematopoietic transition (EHT) whereby they round up and bud into the lumen with concomitant proliferation and thereby decrease endothelial marker gene expression ^17–22^. During this process, the cells undergoing EHT start to gain the hematopoietic markers *cKIT* and CD41 followed by CD45 in a small minority of IAHC, which categories HSCs in the AGM (^23–25^. Moreover, more stringent markers, including SCA1, EPCR can be added to enrich the IAHC for HSCs (T2-HSCs) ^26, 27^. Numerous recent publications have analysed the stepwise progression of HE cells to HSPC at different developmental stages by single cell RNA sequencing ^16, 18, 28–30^. Remarkably, many of them identified the expression of Notch signalling molecules and Notch targets as characteristic for arterial endothelium, HE and IAHC, indicating an essential role in these cell populations ^18, 29^.

The Notch signalling pathway is highly conserved in metazoan and controls cell fate decisions during embryonic development that is initiated by cell-cell contact ^31^. In mammals, there are four Notch receptors (*Notch1–4*) and five ligands: three Delta ligands (*Dll1*, *Dll3*, and *Dll4*) and two Jagged ligands (*Jag1* and *Jag2*) ^32^. Typically, a cell displays one of the Notch receptors and its neighboring cell interacts through a Notch ligand which mediates Notch activation in the receptor bearing cell, thereby releasing the Notch Intracellular Domain (NICD), which then translocate into the nucleus to activate the transcriptional repressor genes, *Hes/Hey* in vertebrates which in turn can repress genes driving cell specification, cell differentiation, and cell cycle arrest ^33^.

Most studies have described Notch receptor and ligands interactions between neighbouring cells (*trans* interactions) that ultimately leads to cell fate segregation within a population by lateral induction or inhibition ^34, 35^. However, emerging studies postulate and demonstrate that receptor and ligand can be co-expressed by the same cell (in *cis*)^36–39^. This *cis* conformation has two advantages for the cell. Firstly, it can switch between different cell fates independent of *trans* activation through expressing *cis*-ligands, and secondly, it can shield or reduce or prevent *trans*-activation by sequestering free receptors. Adding to the complexity of Notch signalling the affinity of the Notch receptors to the ligands can by either weakened or enhanced by glycosylation of the extracellular part of the receptor by fringe proteins (MANIC, LUNATIC and RADICAL) ^40^. Whilst both LUNATIC and MANIC enhance binding of DLL1 to NOTCH1, RADICAL (*Rfng*) also enhances binding through JAG1 ^41^ and, facilitates *cis* activity of NOTCH1 for JAG1^36^.

Since Notch signalling is crucial for both arterial specification and IAHC (including HSC activity), it has been challenging to uncouple Notch signaling requirements for arterial identity from hematopoietic commitment ^42^. In embryonic *Notch1*-chimeras, *Notch1*-deficient cells fail to contribute to haematopoiesis after E15.5, suggesting that *Notch1* is needed in hematopoietic cell intrinsically for their specification ^43, 44^. Counterintuitively, Notch activity tracing transgenic mouse models defined lower Notch activity in IAHC compared to their arterial surrounding ^45^ and blocking DLL4 with a specific antibody at the time of HSC emergence increases HSC frequency ^46^. However, both complete and endothelial specific KO of the notch ligand *Jag1* shows a specific loss of IAHC and HSC activity, although the artery formation is intact ^47, 48^. Finally, maturing T2-HSCs are Notch independent ^49^. It is therefore unclear how these seemingly contradictive phenotypes can be consolidated. Specifically, it is not known how lower Notch activity during HSC maturation through *Notch1* is established.

Here, we studied the dynamic protein levels of notch receptors and ligands by FACS during HE to IAHC transition. We find profound differences in the distribution of the ligands DLL4 and JAG1. DLL4 is predominantly present at early stages and more specifically in HE, whereas JAG1 is detected in robustly in IAHC and its levels are sustained from E10.25 to E11.5. By capturing the surface expression of Notch signaling molecules, NOTCH1, DLL4 and JAG1 at the time of FACS assisted purification of cells (Index FACS) we were able to link their protein expression profile to hematopoietic fate by single cell RNA seq. Remarkably, we identified a subset of Gfi1+ HE that expresses high levels of Notch target genes and suggesting high Notch activity. This small population of Notch activated HE cells proceed to cluster towards T2-HSCs and already express markers of HSC fate. We visualized the interaction of NOTCH1 and DLL4/JAG1 by Proximity ligation assay. We found NOTCH1 and JAG1 interactions accumulated as foci in IAHC, and these interactions could be further identified as *cis* conformation. Furthermore, our single cell RNA seq data set indicated the expression of *Rfng* in a subset of T2-HSCs. By Immunohistochemistry (IHC) and FACS analysis we show that RFNG is specific for the T2-HSC population. Consequently, RFNG knockdown AGMs have reduced numbers of HSCs and the NOTCH1-JAG1 *cis* interactions are reduced, demonstrating that RFNG favors NOTCH1-JAG1 in *cis* and that this conformation is essential for HSC maintenance. Finally, we deregulated this NOTCH1-JAG1 cis interaction by culturing the cells *ex vivo* in the presence of recombinant JAG1 protein (Fc-JAG1). Using a nascent RNA capture assay, we found that genes associated with lineage differentiation and cell cycle are the main networks regulated by NOTCH1-JAG1 in *cis* conformation in Gfi1+ IAHC.

Altogether, we provide here a novel aspect of the role of Notch signalling in HSC biology where RFNG in HSCs facilitates NOTCH1-JAG1 interactions in *cis* that fine tunes the differentiation and cell cycle kinetics to a stem cell state.

## Results

### Notch receptors and ligand proteins are dynamically expressed in the AGM subpopulations

Availability of antibodies against different members of now the Notch family allows the precise characterization of protein levels in rare populations at a single cell level. We characterized the presence of Notch receptors and ligands in the AGM subpopulations during hematopoietic development (E10.25-E11.5). We detected the presence of most of the ligands and receptors at E10.5 and a decrease at E11.5 was observed for NOTCH1 and DLL4 in the endothelium (CD31+cKIT-CD45-) and IAHC (CD31+cKIT+), as well as for JAG1 in CD45+IAHC (Suppl Figures S1and Suppl table T1). Based on the literature and our previous work ^44, 47, 49, 50^, we focused our subsequent analysis on NOTCH1, NOTCH2 receptors and JAG1 and DLL4 ligands.

In order to identify the HE and to restrict the HSC-containing IAHC, we used the *Gfi1:tomato* mouse line (Figure 1A, B). The GFI1 reporter marks a sub-population of the CD31+cKIT-fraction that consists of HE, and a subpopulation of IAHC cells that contain all HSC activity ^11^ (Figure 1B). On average, a large majority of cells in the HE subpopulation (80%) only express DLL4 on their surface, and about 15-20% are NOTCH1+ cells (Figure 1C and D; Suppl Figure S3A). Expression of DLL4 in HE and GFI1+IAHC steadily decreases from E10.5 to E11.5 (Figure 1D, Suppl Fig S2 and S3A), and NOTCH1 expression in GFI1+IAHC suddenly decreases at late AGM stages (Figure 1E and F, Suppl Fig S2 and S3A). In stark contrast, JAG1 protein levels are low in HE (Figure 1 C and D, Suppl Fig S1 and S2A-B), but is the most abundant ligand in GFI1+IAHC (Figure 1 E and F, Suppl Fig S1 and S2A-B). In fact, T2-HSCs, defined as CD31+cKIT+GFI1+CD45+ (Figure 1 A) (^51^), maintain NOTCH1 and JAG1 co-expression on the surface as opposed to CD45 negative CD31+cKIT+GFI1+ subpopulation which loses this co-expression over the course of E10.25 to E11.5 (Figure 1 G and H, Suppl Fig S3C).

**Figure 1:**
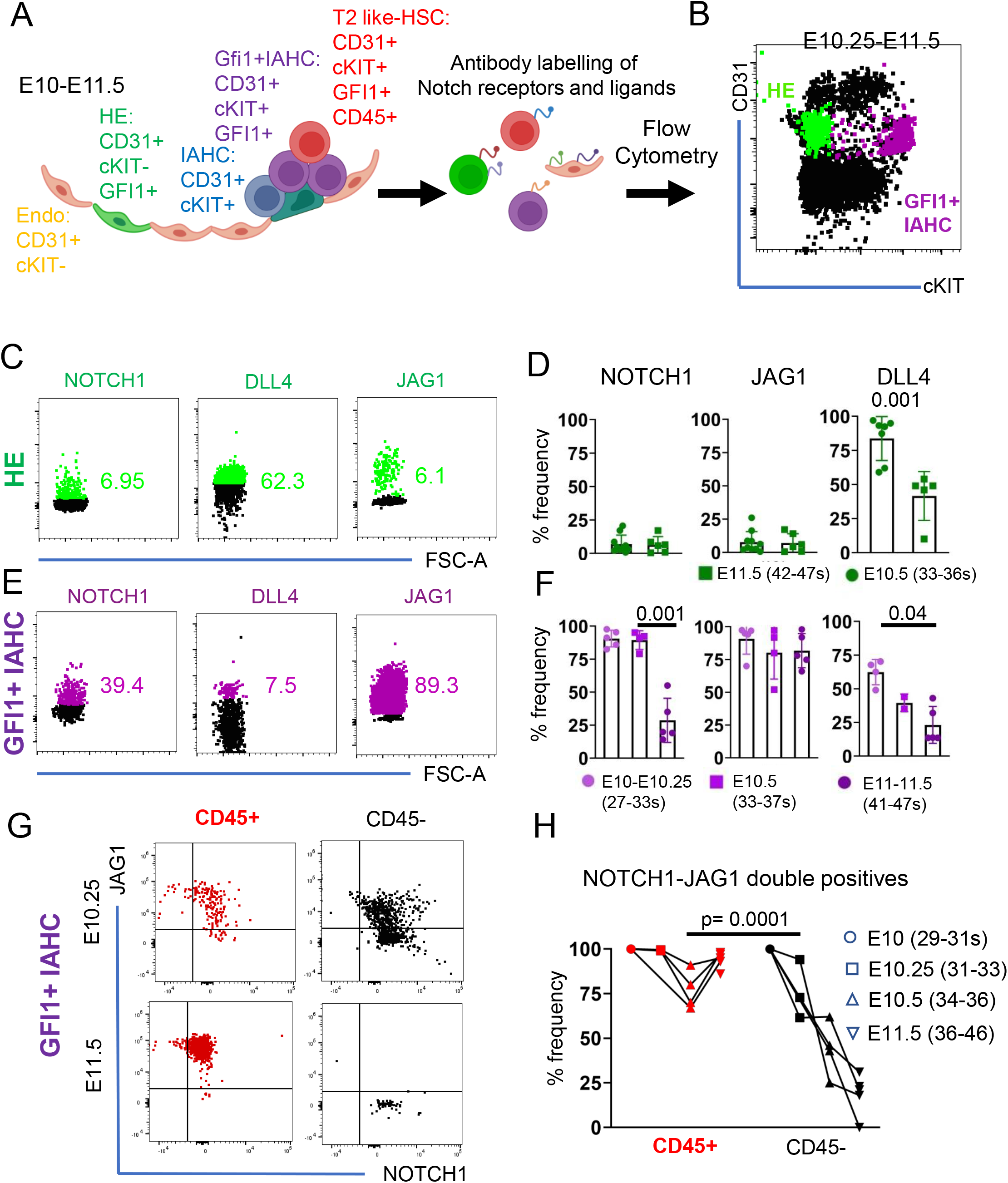
NOTCH1 and JAG1 are co-expressed in T2-HSCs. (**A**) scheme of the IAHC formation in the mouse embryo. GFI1+ HE cells undergo EHT and form IAHC within the dorsal aorta. Within these clusters, the GFI1+IAHC fraction contains all HSC activity. The HSC population can be further restricted by including CD45 as a marker of T2-HSCs (T2-like HSC). (**B**) Representative Flow cytometry plot of AGM lysates stained for CD31 and cKIT. The gate for GFI1+ cells within each population is superimposed onto the plot. (**C**) Representative Flow Cytometry plots of HE with superimposed gates for indicated notch signaling molecule. (**D**) Quantification of NOTCH1, JAG1 and DLL4 positive cells within the HE population at E10.5 and E11.5. 2-5 AGMs were pooled for each data point with a minimum of 4 independent experiments (n ≥4). Statistical significance was calculated with two-tailed t-tests. (**E**) Exemplary Flow Cytometry plots of GFI1+IAHC with superimposed gates for indicated notch signaling molecule. (**F**) Quantification of NOTCH1, JAG1 and DLL4 positive cells within the GFI1+IAHC population at E10.5 and E11.5. 2-5 AGMs were pooled for each data point with a minimum of 4 independent experiments (n≥4). Statistical significance was calculated with two-tailed t-tests. Statistical significance was calculated with two-tailed t-tests. (**G**) Representative Flow Cytometry plots showing JAG1 and NOTCH1 levels within GFI1+ IAHC (CD31+cKIT+GFI1+) sub gated for CD45 at E10.25 and E11.5. NOTCH1-JAG1 double positive cells in the CD45+ fraction is highlighted in red. (**H**) Line chart of Notch1/JAG1 double positive cells within CD45+/- IAHC (CD31+cKIT+) between E10 and E11.5. 2-5 AGMs were pooled for each data point with a minimum of 3 independent experiments for each time point (n>3). Statistical significance was calculated with two-tailed t-tests. Statistical significance was calculated with two-tailed t-tests.

### Single cell RNA sequencing identifies different subpopulations of GFI1+ HE

To further understand the identity of Notch receptor and ligand expressing AGM cells and their trajectory, we performed Index sorting (whereby the FACS data of every sorted cell is recorded) with a panel of hematopoietic markers and Notch-related antibodies combined with single cell RNA sequencing using the *Gfi1:tomato* transgenic embryos (Figure 2A). This approach allows us to link the surface expression of specific Notch signalling molecules to a given cell fate. We index sorted GFI1+ HE (CD31+cKIT-GFI1+) (green), HSC containing IAHC (CD31+cKIT+GFI1+) (purple) and other HPCs (CD31+cKIT+GFI1-) (blue) from E10.5 and E11.5 AGMs (Figure 2A, Suppl Figure S4). A UMAP plot showed how cells from the different index sorted populations overlapped to a large extent except for a subset of Gfi1+HE cells which were not mixed at all (Figure 2B, highlighted with green circle). This subset was mainly obtained from both time points (Suppl Figure S5A) and were potentially in G1 phase (Suppl Figure S5B).

**Figure 2:**
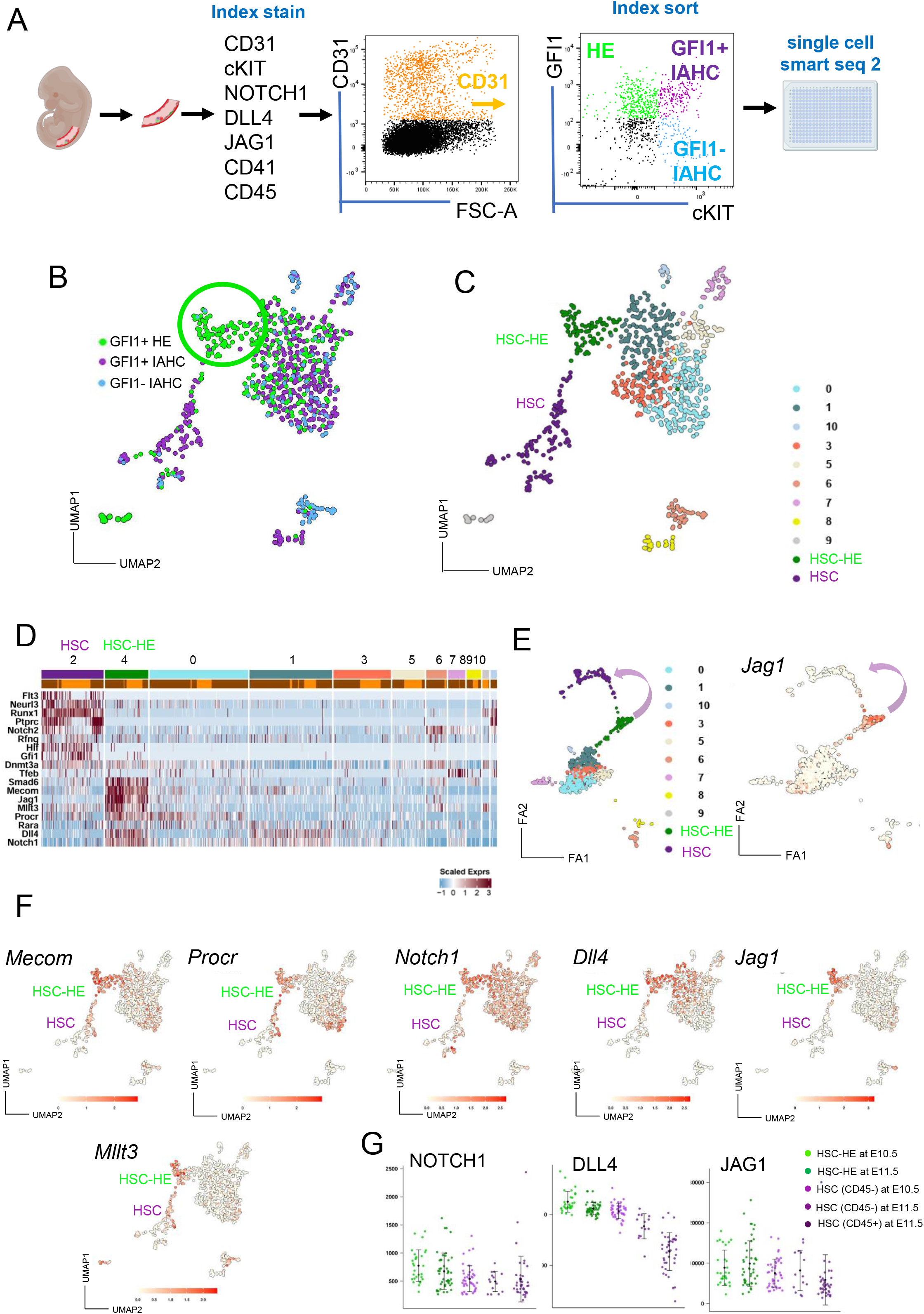
single cell RNA seq of Index sorted HE and IAHC identify HSC-HE and T2-HSC. (**A**) Scheme of the experimental set up. E10.5 and E11.5 AGM lysates were stained for the indicated cell surface markers and Index sorted for Smart seq 2 single cell RNA sequencing. The representative Flow cytometry plot highlights the sorted populations. CD31+GFI1+cKIT-(HE), CD31+GFI1+cKIT+ (GFI1+IAHC) and CD31+GFI1-cKIT+ (GFI1-IAHC). Cells were sorted in 2 independent experiments from a pool of 5 and 6 at E11.5 and 7 at E10.5 AGMs. (**B**) UMAP representation of all Index sorted and sequenced cells. The colors highlight the gate they were sorted with. GFI1+HE (green), GFI1+IAHC (magenta) and Gfi1-IAHC (blue). Green circle highlights the cluster of cells that consists only of cells sorted as GFI1-HE. (**C**) UMAP showing the 11 identified clusters (HSC-HE and HSC are designated). **D**) Heatmap of selected HE and HSC specific genes across all clusters (**E**) Forced directed graph analysis of all sequenced cells with colors highlighting individual clusters and highlighting of *Jag1* levels in forced directed graph analysis. (**F**) UMAP of selected genes with annotation of HSC-HE and HSC. (**G**) Dot plots showing the HSC-HE and HSC at E10.5 and E11.5 with their FACS Index levels for NOTCH1, DLL4 and JAG1. Vertical error bars indicate the mean and standard deviation values.

Clustering analysis of all sequenced cells identified 11 cell clusters (Figure 2C-E) and we obtained the cluster specific marker genes (Suppl Table T2). The top 25 marker genes from clusters 0, 1, 3 and 5 confirmed a high molecular similarity among them, also observed in the UMAP (Figure 2B and C and Suppl Figure S5C). We then assigned cell identity based on already established (^18, 29, 51^) marker gene expression for HE and HSC (Figure 2D) and assigned this phenotype to clusters 4 and 2, respectively (Figure 2C-E).

Next, we assessed the molecular relationship between these clusters by trajectory and pseudo time analysis. We determined that cluster 4 (the pure GFI1+ HE cluster highlighted in green in Figure 2 B) was temporally situated preceding the main two clusters (Figure 2E and Suppl Figure S5D). To our interest, we detected stronger *Jag1* expression levels (Figure 2E) and increasing levels of key HSC markers, including *Mecom, Mllt3* and *Procr* (Figure 2F) from cluster 4 toward the more distal HSC cluster (Figure 2C-F). We therefore will refer to this sub-population of GFI1+ HE as HSC-primed HE (HSC-HE). Within the distal HSC cluster (cluster 2, mostly composed of GFI1+IAHC cells) (Figure 2B), we detected T2 HSC associated marker gene expression (Figure 2D; Suppl Figure S5E) and *Flt3*, a newly identified marker for embryonic multi-potent progenitors (Suppl Figure S5F)^8^.

Subsequently, we analysed the specific Notch distribution in these AGM subpopulations. We found that *Notch1*, *Dll4* and *Jag1* gene expression was highly abundant in the HSC-HE clusters with *Jag1* being distinct for this cluster (Figure 2E and F), suggesting that this ligand has an important role in HSC biology. Taking advantage of the index FACS data, we further confirmed that JAG1 and NOTCH1 proteins were also present at the surface of the cells in the HSC-HE and HSC clusters (Figure 2G; Suppl Figure S5G).

### High *Notch* transcriptional activity identifies a HSC-primed HE cluster and decreases with HSPC maturation

To understand the Notch activity dynamics, we plotted the expression of direct Notch targets across the UMAP (Figure 3A green circle highlighting the HSC-HE). We observed heterogeneous *Hes1* and *Gata2* expression among the different clusters, but *Hey1*, and specially *Hey2,* were mainly restricted to the HSC-HE cluster (Figure 3A, HSC-HE highlighted in green). We therefore examined the molecular differences between GFI1+ HE that were co-expressing *Hey1/2* (HSC-HE) and compared it to the rest of GFI1+ HE at the different time points. Besides expressing *Hey1* and *Hey2*, we observed an upregulation of *Notch1* and *Jag1* in both time points (Figure 3B-D). Differential gene expression analysis (DEA) indicated that several HSC-associated genes were exclusively up regulated in the *Hey1/2* positive (HSC-) HE at E10.5 and E11.5 (Figure 3B-D, Suppl table T3). KEGG pathway analysis further enriched gene sets that are known for their involvement in HSC emergence, i.e., signalling pathways in pluripotency in stem cells, HIF1, shear stress, and TNF–signalling (Figure 3E, Suppl table T3). Interestingly, Notch signalling pathway was significantly overrepresented at both E10.5 and E11.5 time points in the *Hey1/2* positive (HSC-) HE cluster (Figure 3E, suppl table T3).

**Figure 3:**
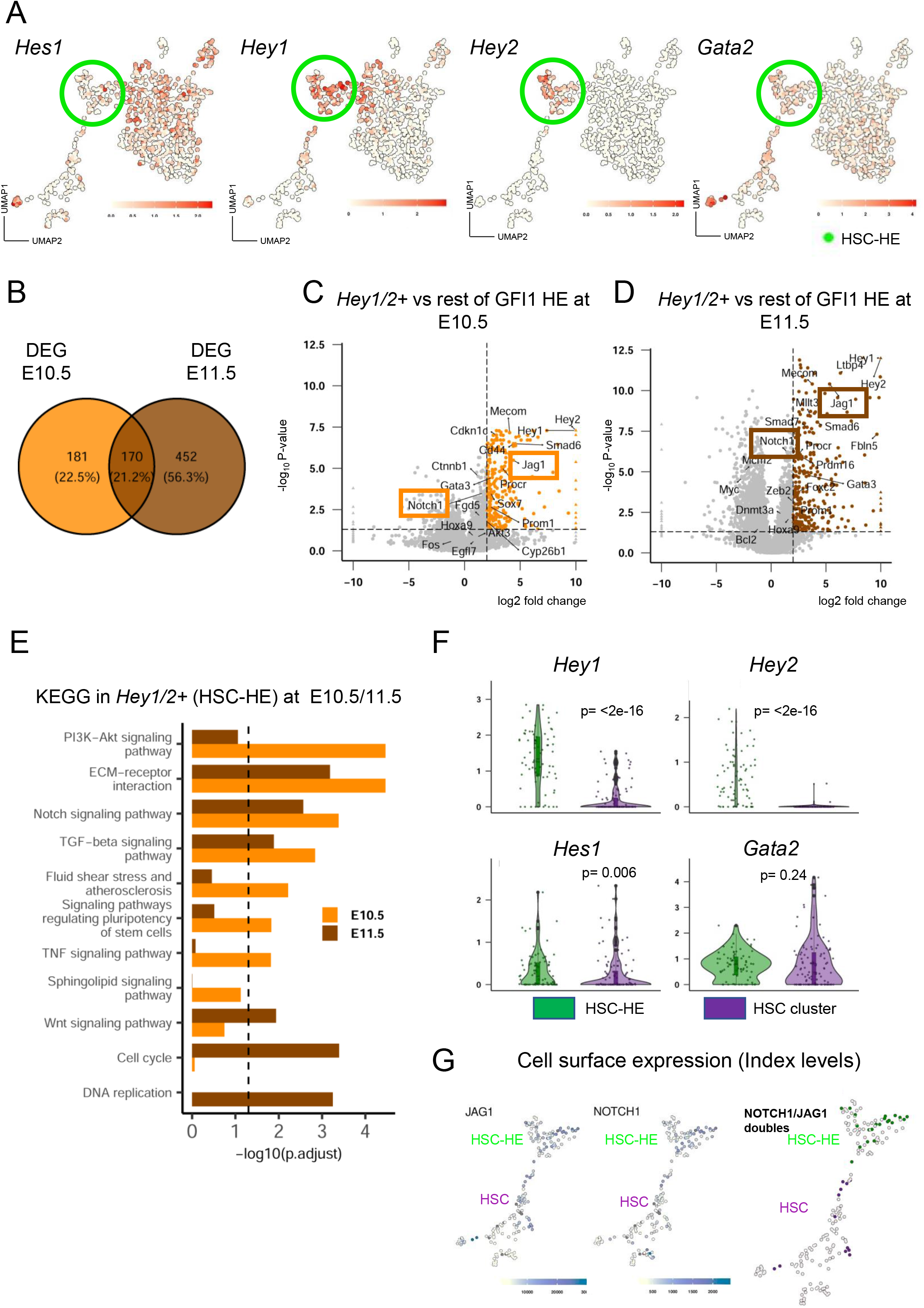
*Hey1/2* expression in HE marks the onset of the HSC specific gene program. (**A**) UMAP representation of *Hes1*, *Hey1*, *Hey2* and *Gata2* expression levels with green circle highlighting the HSC-HE population. (**B**) Venn diagram of DEGs in GFI1+HE cells expressing *Hey1/2* compared to the rest at E10.5 and E11.5. (**C**) and (**D**) Volcano plots of DEGs between *Hey1/2+*HE compared to the rest of the GFI1+HE at E10.5 (351 genes) and at E11.5 (622 genes) respectively. Genes that are upregulated above log_2_ Fold Change > 1 and adjusted p-value (FDR) < 0.05 are highlighted in orange (E10.5) or brown (E11.5). *Jag1* and *Notch1* expression are pointed out with a box. Genes showing absolute log_2_ Fold Change > 10 are plotted as triangle (**E**) KEGG pathways overrepresentation analysis r of DEGs of *Hey1/2+*HE compared to the rest of the GFI1+HE at E10.5 (orange) and E11.5 (brown). (**F**) Violin and boxplots of gene expression levels of *Hes1*, *Hey1*, *Hey2* and *Gata2* comparing HSC-HE and T2-HSCs. Boxplots show the median (centre line) first and third quartiles (box limits), and a maximum of 1.5x the interquartile range (whiskers). Statistical significance was calculated with Wilcoxon test (**G**) UMAP representation of NOTCH1 and JAG1 co-expressing cells (protein) derived from the index label with HSC-HE and HSC.

To determine Notch activity dynamics, we compared Notch-target gene expression in the HSC-HE and the HSC cluster. We observed a significant downregulation of *Hey1*, *Hey2* and *Hes1*, while *Gata2* expression was maintained (Figure 3F). The persistence of the JAG1 and NOTCH1 protein from HE to HSCs (Figure 1G/H and 2G) but the gradual decrease of Notch target genes from the HE to T2-HSCs (Figure 3A and 3F) led us to the hypothesis that JAG1 was not participating in Notch receptor activation (where it would be rapidly endocytosed), but in an alternate function.

### JAG1 and NOTCH1 interactions accumulate in IAHC

Next, we performed IHC to validate our FACS Index distribution of JAG1 and DLL4 in GFI1+ AGM sections. Indeed, we detected both DLL4 and JAG1 in GFI1+ IAHC clusters. Interestingly, we discovered differences in the spatial distribution of these two ligands (Suppl Fig S6A). DLL4 was present as discreet foci, whereas JAG1 expressed diffuse across the whole cell surface of GFI1+ cells (Suppl Figure S6A). This observation prompted us to study the interactions between the NOTCH1 and the two ligands DLL4 and JAG1 using Proximity Ligation Assay (PLA). In this assay, each point of interaction between a receptor and its ligand is visualized as a fluorescent foci/signal. Here, DLL4, NOTCH1 and JAG1 antibodies are labelled with oligonucleotides and can serve as primers for rolling circle amplification only if they are close enough to interact (<30nm). We multiplexed the PLA for two sets of antibodies (and compatible oligos). One set of PLA probes/antibodies target the extracellular (N-terminal) domains of the NOTCH1 (NOTCH1-extra) and DLL4 (DLL4-extra) and produces a signal in yellow (Figure 4A, yellow). The second pair of antibodies/probes binds the extracellular (N-terminal) domains of the NOTCH1 (NOTCH1-extra) and JAG1 (JAG1-extra) with interactions producing a fluorescent signal in far red (Figure 4A, magenta). These two sets were used to probe thick (150um) AGM sections of E10.5-E11.5 embryos and imaged as serial confocal images (z-stacks). The 3D rendering of the z-stacks shows punctuated staining for both NOTCH1-extra/DLL4-extra (yellow) and NOTCH1-extra/JAG1-extra (magenta) as expected for this type of assay (Suppl Figure S6B and C). For NOTCH1-extra/DLL4-extra interactions (yellow), we saw a low distribution of fluorescence in endothelial cells and the surrounding tissue, with increased dots also detected in IAHC (Figure 4B, left and right). Similarly, we detected sparse NOTCH1-extra/JAG1-extra (magenta) in all AGM tissues, but a much clearer and greater level of signal than for NOTCH1-extra/DLL4-extra in IAHC (Figure 4B, middle and right). We quantified the total number of amplification dots for each PLA pair in endothelial cells (control) and compared it to the number of dots detected in IAHC. We did not see any significant difference in the fluorescence accumulation for NOTCH1-extra/DLL4-extra (yellow) or NOTCH1-extra/JAG1-extra (magenta) in endothelial cells, but we established significantly higher levels of NOTCH1-extra/JAG1-extra than NOTCH1-extra/DLL4-extra in IAHC (Figure 4C).

**Figure 4:**
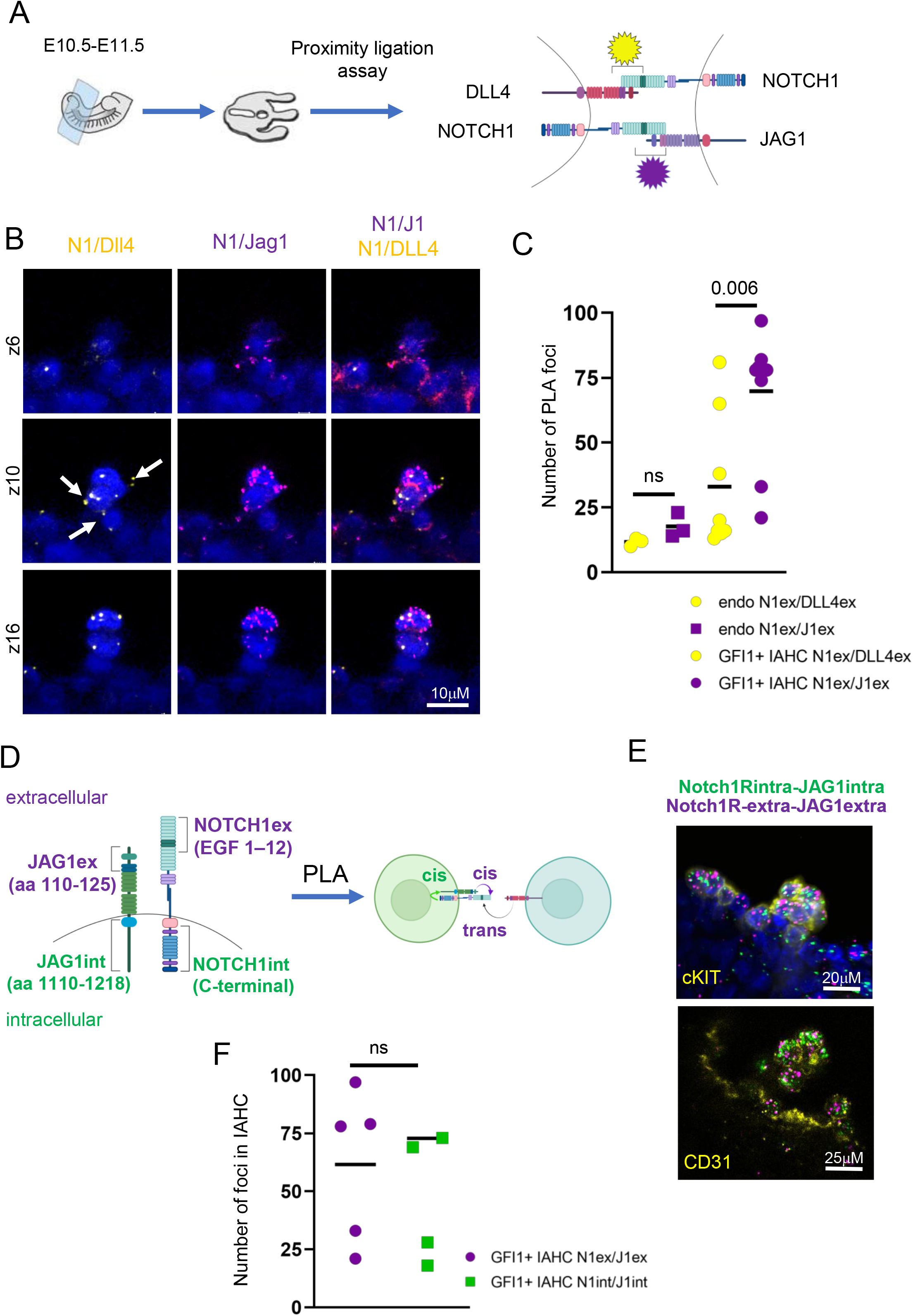
NOTCH1 and JAG1 for a cis interaction in IAHC. (**A**) Scheme of experimental set up. 150μm thick trunk sections of E11.5 AGMs were subject to Proximity ligation assay (PLA) with indicated antibody pairs followed by confocal imaging. (**B**) Representative optical 2-3 μm sections (z-stacks,z) through a IAHC. PLA signals are detected as spots for points of interactions. NOTCH1/DLL4 (yellow), NOTCH1ext/JAG1ext (magenta) and DAPI. Scale = 10μm. (**C**) Quantification of foci (interaction points) for NOTCH1/DLL4 (yellow), NOTCH1ext/JAG1ext (magenta) in endothelial cells (endo) or IAHC. Statistical significance was calculated with two-tailed t-tests (n= 4 independent experiments). (**D**) Scheme of experimental set up to distinguish between *trans* and *cis* NOTCH1-JAG1 interactions with antibodies recognizing the indicated amino acid sequence of NOTCH1 or JAG1. NOTCH1ext/JAG1ext (magenta) and NOTCH1int/JAG1int (green). (**E**) Exemplary images of a cKIT+ or CD31+ IAHC probed with PLA with NOTCH1ext/JAG1ext (magenta) and NOTCH1int/JAG1int (green) and DAPI. (**F**) Quantification of foci (interaction points) for NOTCH1ext/JAG1ext (magenta) and NOTCH1int/JAG1int (green) in IAHC. Statistical significance was calculated with two-tailed t-tests (n= 3 independent experiments).

To validate the specificity of the PLA for Notch interactions, we did a control experiment with cells overexpressing *Manic fringe* (MFNG). MFNG is a glycosyltransferase that enhances the binding of DLL4 to NOTCH1. We obtained *Tie2:Mfng* overexpressing AGMs and performed the PLA for both antibody pairs. As expected, and further confirming the validity of the assay, we detected several enhanced interactions for NOTCH1-extra/DLL4-extra (yellow) in IAHC whereas NOTCH1-extra/JAG1-extra (magenta) did not change (Suppl Figure S6D).

Curiously, and in agreement with our IHC staining (Suppl Figure S6A), most interactions between NOTCH1-extra/DLL4-extra (yellow) in IAHC were located between cells (Figure 4B white arrows and Suppl Figure S6C white arrows), indicating a possible *trans* interaction between adjacent cells. On the contrary, NOTCH1-extra/JAG1-extra (magenta) interactions were frequently detected as interactions that cover the entirety of the cell surface in IAHC, even in the absence of neighboring cells (Figure 4B, middle and right and Suppl Figure S6C). Consistent with FACS and index sorting data that shows NOTCH1 and JAG1 on the surface of the same cell (Figure 1H, 2F and G, 3G), and the interaction pattern of NOTCH1 and JAG1 by PLA, raised the possibility that NOTCH1 and JAG1 might be interacting on the surface of the same cell (*cis*). This mode of interaction might shield the IAHC from further NOTCH1 interaction with ligands presented from the surrounding cells.

### JAG1 and NOTCH1 interact in *cis* in IAHC

To test this hypothesis, we probed the intracellular domains (C-terminus) of the NOTCH1 and JAG1 with the rationale that we would only detect a signal if these intracellular parts were a) on the same cell, i.e. in parallel orientation to each other and b) therefore close enough to interact, i.e. be in a *cis* configuration (Figure 4D). We multiplexed the new PLA pair, NOTCH1-int/JAG1-int (green) with the two previous probe pairs and analysed their fluorescence distribution in the AGM sections. Most importantly, we quantified the number of interactions from NOTCH1-extra/JAG1-extra (magenta) and NOTCH1-int/JAG1-int (green) in IAHC. In agreement with our hypothesis, we detected accumulation of the green NOTCH1-int/JAG1-int signal specifically in cKIT+ cells/ IAHC (Figure 4E; suppl Figure S6E). Intriguingly, we did not find a significant difference in the number of interactions for NOTCH1-extra/JAG1-extra (magenta) and NOTCH1-int/JAG1-int (green) (Figure 4F), implying that most NOTCH1 interactions with JAG1 were detected targeting both the intracellular domains and the extracellular domains. These results strongly suggest that *cis* interactions between NOTCH1-JAG1 (detected by both NOTCH1-int/JAG1-int and NOTCH1-extra/JAG1-extra) are occurring in IAHC since no increase is detected in NOTCH1-extra/JAG1-extra interactions (it should detect both *cis* and *trans*).

Consistent with the proposed role of *cis* interactions in the Notch pathway, they have the potential to interfere with Notch activity and could be the cause for the gradual decrease observed from HSC-HE to T2-HSC for NOTCH target gene expression.

### *Cis* interaction between JAG1 and NOTCH1 reduces Notch activation in IAHC

This intriguing new finding prompted us to speculate that if the role of JAG1 in IAHC was to shield them from Notch signaling activation, then disrupting this interaction could consequently lead to alterations in Notch activity. We therefore purified endothelial cells (CD31+cKIT-GFI1-) as a reference and IAHC (CD31+cKIT+GFI1+) in parallel from E11.5 AGMs (Figure 5A). Both populations of cells were incubated overnight with the γ-secretase inhibitor, compound E (CompE), to abolish basal Notch activity. Both populations were subject to different conditions for 4 hours after which the cells were harvested for gene expression analysis (Figure 5A). The cells were either kept in a Notch inhibited state (compE), released out of Notch inhibition (wash), or stimulated with soluble, recombinant JAG1 (Fc-JAG1) (Figure 5A), being our prediction that excess of Fc-JAG1 would activate Notch signaling in *trans* and disrupt *cis*-inhibition. To minimize cell-cell interactions, cells were seeded at low density (300-400 cells/well) in semi-solid methylcellulose (with growth factors). We supplemented the methylcellulose culture with a nascent RNA capture reagent (EU) to capture and quantify the transcriptional changes that occur in each condition and population. We first validated this experimental approach by performing qPCRs for the Notch targets *Hes1* and *Gata2* since they are both expressed in endothelial cells and HSCs (Figure 3A). We detected lower levels of both in the compE condition and higher levels with Fc-JAG1 stimulation (Figure 5B), confirming that excess of Fc-JAG1 was activating NOTCH1. However, the up-regulation of *Hes1* and *Gata2* in response to Fc-JAG1 was more robust in endothelial cells than in GFI1+IAHC cells (Figure 5B), indicating that GFI1+IAHC cells are more refractory to Notch activation. These results are compatible with a *cis*-inhibitory function of JAG1 in GFI1+IAHC cells.

**Figure 5:**
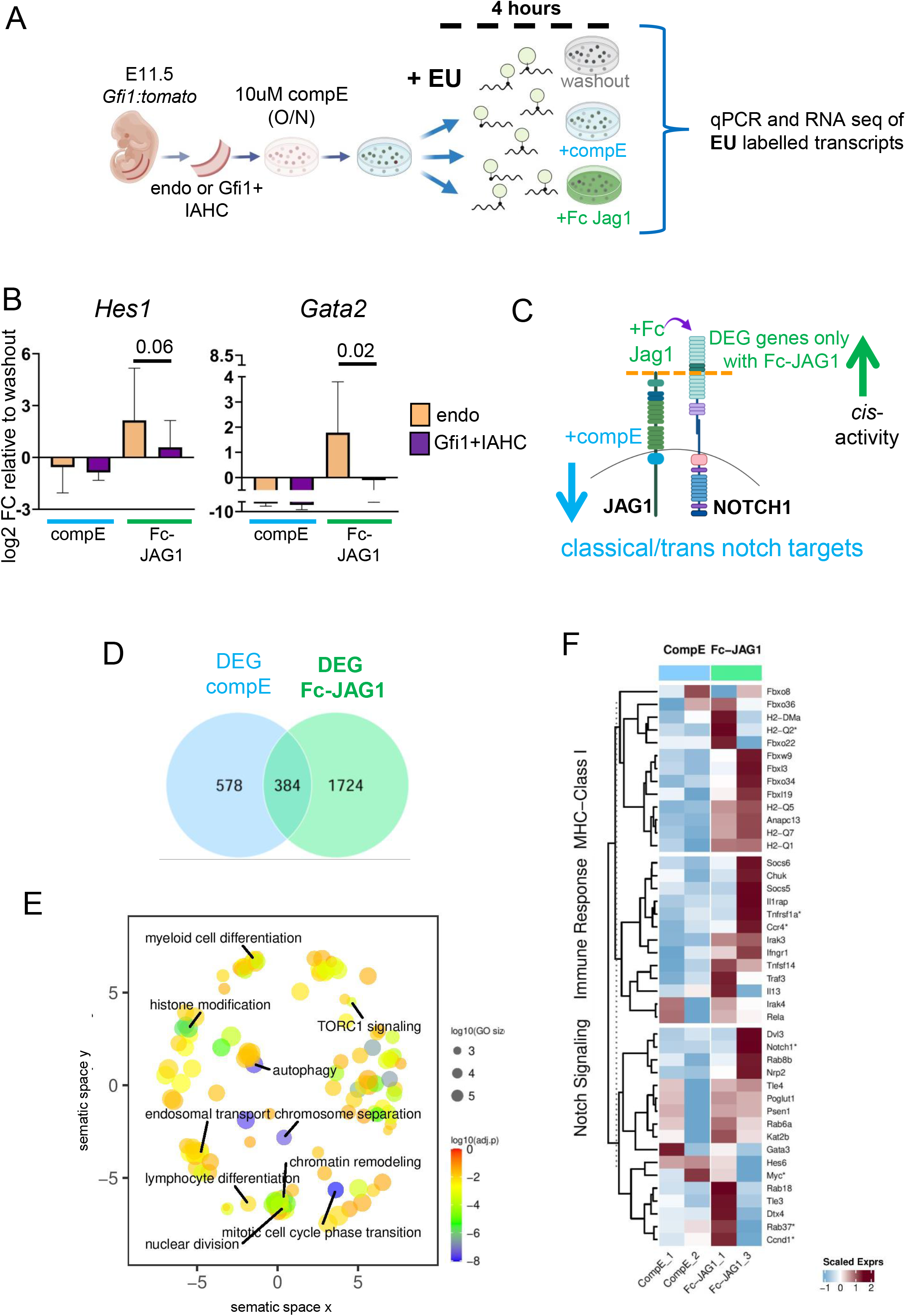
JAG1 in GFI1+IAHC controls stemness related pathways. (**A**) Scheme of experimental procedure. Endo (CD31-GFI1-ckit-) and GFI1+IAHC from E11.5 AGM were FACS purified and treated with 10μM of compE overnight. The cells were then washed and split into 3 conditions as illustrated. Either cells remained in a washout condition (control), returned to 10μM of CompE treatment (CompE) or cultured with Fc-JAG1. 3 independent experiments with cells purified from pools of embryos (at least 5 E11.5 AGM per experiment) were performed. All conditions were cultured in semi-solid methylcellulose at low concentration to limit cell-cell contact and 10μM of EU was added to each culture to label the newly synthesized RNA for 4 hours. After this time, RNA was extracted, cDNA was pre-amplified and subject to qPCRs and RNA sequencing. (**B**) Relative gene expression levels of *Hes1* and *Gata2* in compE and Fc-JAG1 treated samples compared to washout control by qPCR. Results were obtained from 3 independent replicates and statistical significance was calculated with two-tailed t-tests. (**C**) Illustration of NOTCH1 and JAG1 in a *cis* conformation of a cell and the expected deregulation of notch targets in response to compE treatment (blocks all Notch activity) and Fc-JAG1 (binds to freely available NOTCH1 or activates NOTCH1 by competing with the JAG1 in *cis*). (**D**) Venn Diagram of DEGs (adjusted p-value (FDR) <0.05) obtained from the independent comparison of CompE and Fc-JAG1 conditions replicates against the washout condition, considered as the control group. (**E**) Gene ontology enrichment of genes that are only significantly over-represented in samples treated with Fc-JAG1. (**F**) Heat map of representative genes that belong to the indicated gene classification and their expression level in CompE and Fc-JAG1 treated samples. Only genes that are differentially expressed between CompE and Fc-JAG with a p<0.05 are considered.

### NOTCH1-JAG1 interactions in *cis* inhibit lineage differentiation genes

To further understand the effects of Notch manipulation in nascent transcription, we proceeded to sequencing the nascent RNA of the Gfi1+ IAHC samples after Notch signaling manipulation (Suppl Figure S6F). Differentially expressed genes (DEGs) were independently identified for compE and Fc-JAG1 conditions relative to the washout, which was considered as the reference (Figure 5C). Those DEGs in the compE scenarios (962 genes) were classified as classical Notch responders (Figure 5D, Suppl table T4). Part of them (384) were in common with those obtained from Fc-JAG1 comparison. Finally, a total of 1,024 genes were exclusively differentially expressed when Fc-JAG1 was added to the culture. From these, the set of up-regulated genes encompassed genes associated to mitotic cell cycle transition, myeloid and lymphoid cell differentiation, chromatin remodeling and autophagy (Figure 5E, Suppl table T4). The enrichment of these Gene ontology terms suggests that disrupting the *cis* inhibition (NOTCH1-JAG1) by *trans* activation through Fc-JAG1 triggers cell cycle entry and activation of lineage differentiation program. We further identified a more naïve state of the GFI1+ IAHC before incubation with Fc-JAG1 by examining genes involved in MHC class I antigen presentation/processing and Immune response since naïve HSCs typically show lower abundance of these group of genes^52, 53^. We found several genes of these two processes significantly up-regulated upon Fc-JAG1 stimulation (Figure 5F and Suppl table T4). Finally, Notch signaling-related molecules were also up-regulated upon stimulation with Fc-JAG1.

In summary, we find evidence that within the GFI1+ IAHC cells that retain NOTCH1 and JAG1 in *cis* conformation, Fc-JAG1 causes activation of genes associated with cell cycle entry and differentiation. These results agree with our hypothesis that the function of the *cis* conformation is to preserve a naïve HSC state within the GFI1+ IAHC cells.

### NOTCH1-JAG1 interactions in *cis* is dependent on RFNG

Notch post-translational modifications by FRINGE glycosyltransferases can enhance ligand-receptor interactions in a specific way. Next to *Mfng* (not shown), *Rfng* expression was also detected in the T2-HSC population (Figure 2D). RFNG has been reported to enhance NOTCH1-JAG1 *cis* interactions in cultured cells^36^. To test whether the *cis* interaction in HSCs could be modulated by RFNG, we performed IHC for RFNG on *Gfi1:tomato* AGMs. We detected RFNG positive cells in a few GFI1+IAHC of E11.5 AGMs (8 out of 19). To quantify the RFNG positive cells within the whole AGM, we performed FACS analysis, combined with the HSC markers, Sca1 and EPCR (T2-HSCs)^51^. We detected significant accumulation of RFNG positive cells in T2-HSCs and some EPCR positive cells, but not in Sca1-EPCR-IAHCs (Figure 6B and C; Suppl Figure S7A). We went further and sub-gated the T2-HSC fraction for RFNG+/- cells to determine if these cells were co-expressing NOTCH1 and JAG1 in the *cis* confirmation. In support of our hypothesis, RFNG positive T2-HSCs enriched for NOTCH1 and JAG1 co-expression (Figure 6D and E).

**Figure 6:**
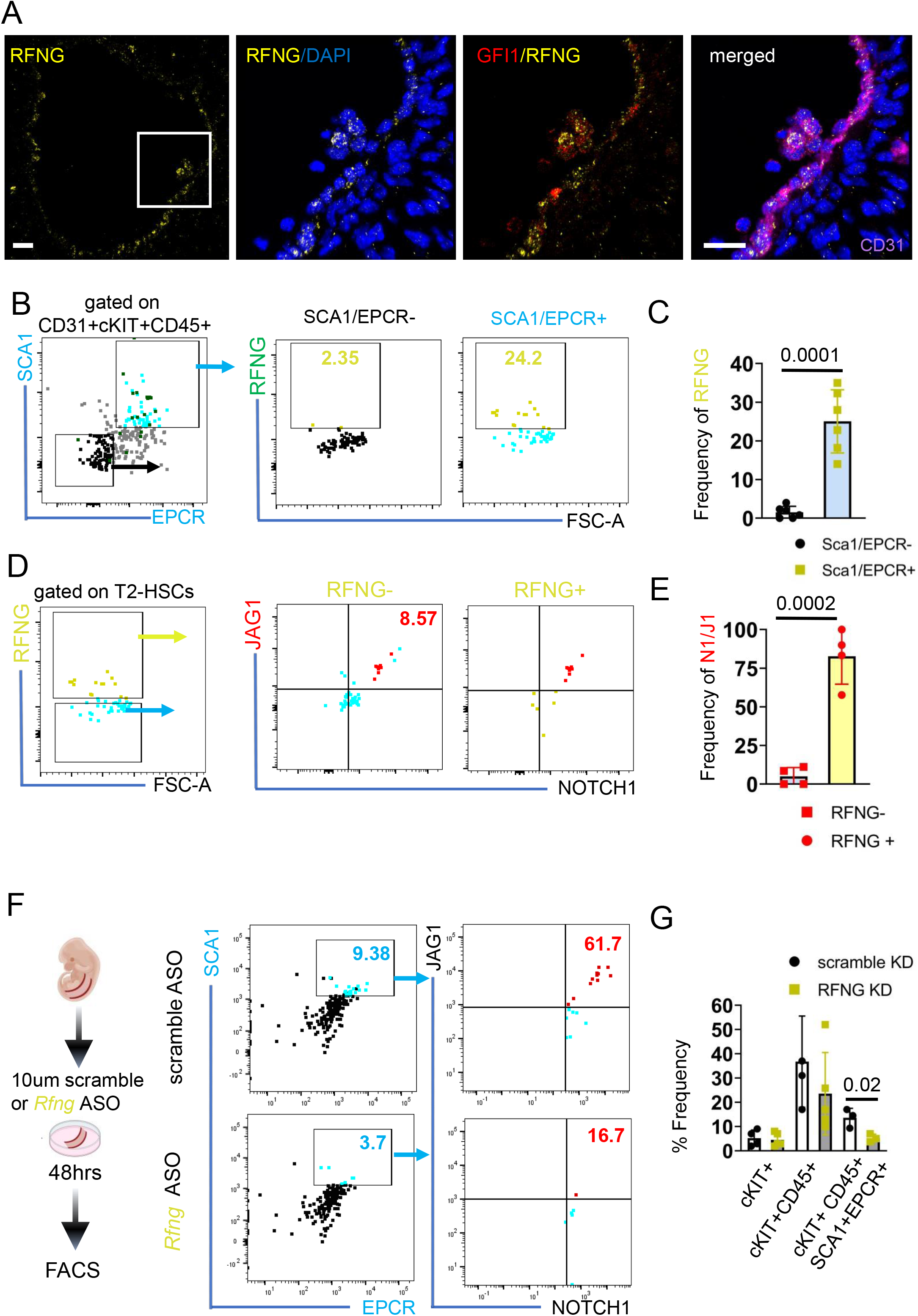
NOTCH1/JAG1 cis interaction and HSC emergence is dependent on RFNG. (**A**) IHC on E11.5 AGM sections for RFNG (yellow), GFI1 (red), CD31 (magenta) and DAPI (blue). Scale bar = 10μm. (**B**) Flow cytometry plots for Sca1 and EPCR after gating for CD31+/cKIT+/CD45+. SCA-/EPCR- or SCA+/EPCR+ and further analysed for RFNG. RFNG gate is superimposed onto Sca1 and EPCR plot. (**C**) Quantification of the frequency of RFNG in either SCA-/EPCR- or SCA+/EPCR+ fraction of CD31+/cKIT+/CD45+. Statistical significance was calculated with two-tailed t-tests. (n=3 independent experiments with 5-10 AGMs each per sample). (**D**) CD31+/cKIT+/CD45+Sca1+EPCR+ T2-HSCs were gated for RFNG expression. RFNG- and RFNG+ T-HSCs were further plotted for JAG1 and NOTCH1. (**E**) Quantification of the frequency of JAG1/NOTCH1 in either RFNG- or RFNG+ fraction of T2-HSCs. Statistical significance was calculated with two-tailed t-tests. (n=2 independent experiments with 5-10 AGMs each per sample) (**F**) Cartoon of experimental set up. E10.5 AGMs were cultured as explant with 10μM control (scramble) FANA-ASO or 10μM of RFNG FANA-ASO and processed for FACS analysis 48hrs later. Flow cytometry plots for Sca1 and EPCR after gating for CD31+/cKIT+/CD45+ for scramble FANA-ASO or RFNG FANA-ASO. The T2-HSCs were then further analysed for JAG1 and NOTCH1 levels. (**G**) Quantification of the frequency of cKIT, cKIT+CD45+ and cKIT+CD45+EPCR+Sca1+ T2-HSCs in either scramble or RFNG ASO. (n=2 independent experiments with 4-5 AGMs each per sample). Statistical significance was calculated with two-tailed t-tests.

**Figure 7:**
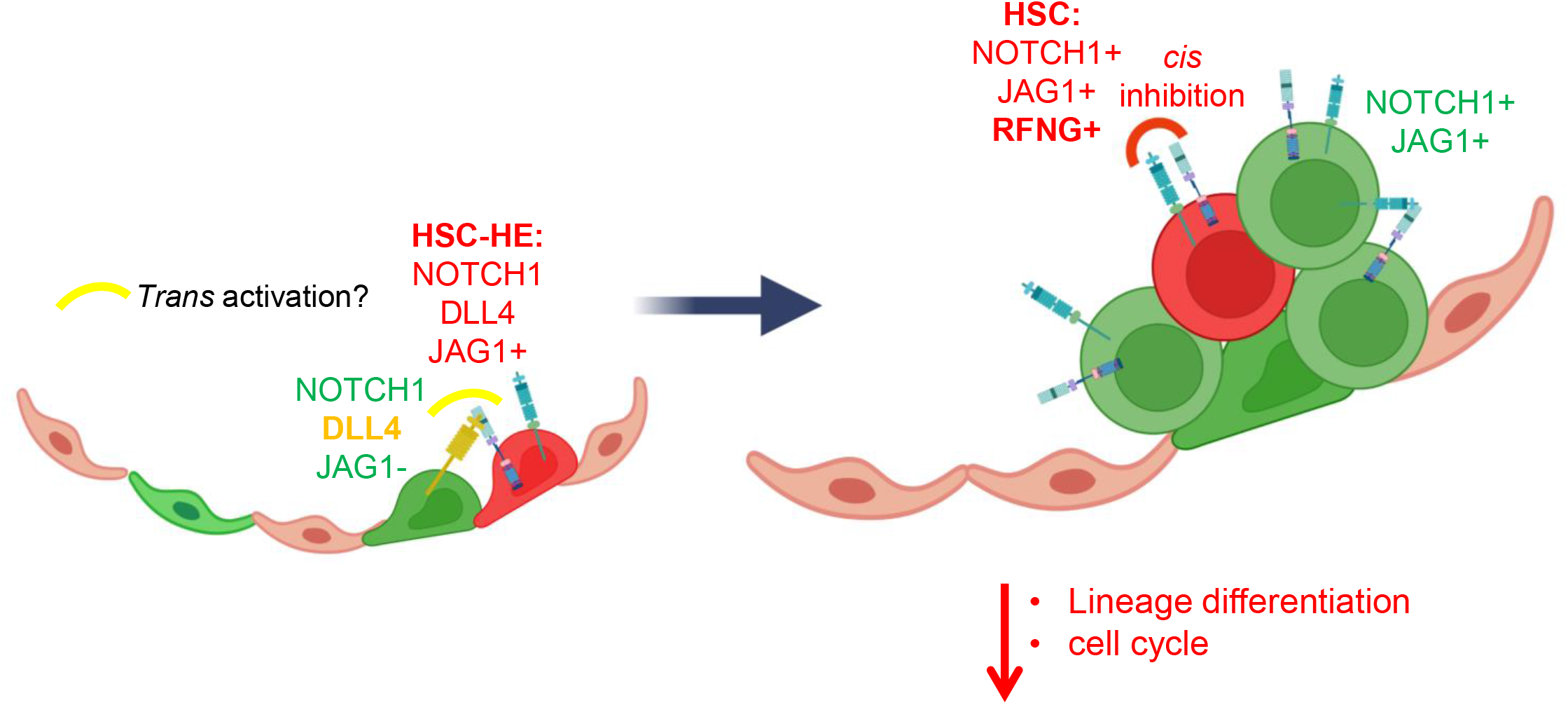
model of Notch signaling in HSC emergence. Scheme of proposed modulation of HSC specification through RFNG. GFI1 HE can be separated into HSC-HE that expresses higher levels of *Jag1*. Within the GFI1+ IAHC, some JAG1 positive cells co-express NOTCH1 on the surface, and this co-expression exists as a *cis* (inhibitory) conformation due to the presence of RFNG. RFNG expression is restricted to the T2-HSC population and this *cis* interaction is necessary to maintain stem cell identity by regulating cell cycle and differentiation associated genes.

Finally, we performed knock down experiments of RFNG in *ex vivo* AGM cultures with anti-*Rfng* FANA-ASO nucleotides (Figure 6F). These nucleotides enter the cell, bind to the transcript and recruit RNaseH to degrade the mRNA (Suppl Figure S7B). We cultured control (scramble FANA-ASO) or *Rfng* ASO treated AGMs for two days and analysed the T2-HSC and NOTCH1-JAG1 co-expression frequency by FACS (Figure 6F; Suppl Figure S7B). In agreement with our hypothesis, a decrease in RFNG in the AGM resulted in significantly fewer phenotypic HSCs (CD45+CKIT+SCA1+EPCR+) concomitant with a reduction in NOTCH1-JAG1 co-expression (Figure 6F-G).

Altogether indicates that RFNG is important to preserve the NOTCH1-JAG1 cis interaction in some IAHC cells and maintain their HSC phenotype.

## Discussion

Notch signalling is required for a plethora of cellular decisions that control cell states and cell fate acquisition, including hematopoiesis. In the embryonic aorta, hemogenic endothelial cells give rise to HSPCs in close association with the aortic endothelium. Arterial and hemogenic endothelium and HSPCs are specified in the AGM region, and all require a specific level of Notch activity. Earlier studies have elucidated that NOTCH1, JAG1, RBPJ and HES repressors are key for the generation of HSPC ^54–57^, while NOTCH1, NOTCH4, DLL4 and RBPJ are required for arterial development ^58–61^. Therefore, it is concluded that Notch activity is compulsory for HSC emergence and arterial formation, but the timing and the threshold of Notch activity as well as the precise interactions of receptors and ligands remained undetermined. In the AGM, blocking DLL4 or inhibiting Notch activity with γ-secretase inhibitors can increase HSPC activity ^62–64^, yet absence of JAG1 leads to a dramatic loss of IAHC ^65, 66^. Importantly, the phenotypes of blocking DLL4 with an antibody or treatment with γ-secretase inhibitors is dependent on the developmental stage of HE and HSPC; blocking DLL4 *ex vivo* most efficiently increases HSC activity when applied at early AGM stages (31-34s) ^62^. Likewise, blocking of Notch activity with γ-secretase inhibitors before acquiring HSC fate can diminish HSC activity ^49^. Taken together, we conclude that Notch activity needs to be dampened once the (HSC)-HE has been specified. In agreement with this assumption, Notch activity tracing lines detect lower notch activation in IAHC than their arterial surrounding ^45, 63, 67, 68^. Here, we present evidence for a *cis* interaction between NOTCH1 and JAG1 that is especially relevant in T2-HSCs and facilitated by RFNG that integrates and further explains the previous observations. Increasing JAG1 levels on the cell surface blocks Notch signalling by forming a *cis* (inhibitory) conformation with NOTCH1. A previous study already established higher levels of activated NOTCH1 (NICD) in IAHC in the absence of JAG1. The *Jag1* deficient cKIT+ IAHCs were present in the aorta but could not bud into the lumen of the aorta, and instead stayed embedded in the endothelium and expressed high levels of endothelial specific genes, indicating that hematopoietic maturation was blocked ^45^. Similarly, and in support for a opposing effects for DLL4 and JAG1 in HSC specification, AGMs treated with a DLL4 blocking antibody show enhanced HSC activity and reduN target gene expression^46^.

The time course of Notch ligand and receptor expression in the hemogenic and IAHC subpopulations suggests that this *cis* interaction is specifically maintained in T2-HSC, while other HPC lose this co-expression. We speculate that a fine balance of Notch activity that is most likely induced by *trans* activation by DLL4 and increasing *cis* inhibition through JAG1 are key for permissive conditions of HSC fate. DLL4 activation should be inhibited since it has a detrimental effect over HSC activity^45^, however *cis* inhibition could also prevent NOTCH1 from responding to free JAG1 or other expressed ligands such as JAG2^55^. We demonstrate that the persistence of this *cis* NOTCH1-JAG1 is dependent on RFNG expression, a glycosyltransferase that modifies the NOTCH1 receptor to have a stronger affinity for JAG1 in *cis* ^36, 69^. Accordingly, we find RFNG concentrated in sparse cells within a few IAHC by IHC and by FACS, we specifically detect them in EPCR+/Sca1+/ T2-HSC. Furthermore, inhibition of *Rfng* results in a reduction of NOTCH1-JAG1 co-expressing T2-HSCs. By nascent RNA capture assay, we identified the pathways that are regulated by JAG1 in the HSC population. We find enrichment of gene sets that include cell cycle, chromatin remodeling, lineage differentiation, and antigen processing/presenting genes that are upregulated in response to Fc-JAG1, strongly indicating that the *cis* confirmation of NOTCH1-JAG1 preserves a naïve stem cell fate phenotype.

It is interesting to note that some HSC specific genes, including *Procr*, *Mecom*, *Fgd5*, and *Hoxa9* are expressed at higher levels in a subset of the GFI1+HE that has high levels of *Jag1* and some Notch target gene expression (Figure 2D-F and 3A). This suggests that the HSC fate is established as early as the HE state, maybe through Notch signaling and this is preserved in the IAHC. In addition, we find that these precursors have a low proliferation index to ensure their integrity (Suppl Fig S5B). Of note, NOTCH1 mutant AGM explants showed significantly higher levels of BrdU+ incorporation in the hematopoietic compartment ^63^ suggestive of Notch signalling controlling cell cycle dynamics in the HE/IAHC. The notion that the HE is heterogeneous in regards of their hematopoietic potential has been recently postulated ^70, 71^. We are therefore further confirming this finding and are proposing a mechanism, namely NOTCH1-JAG1 in *cis*, as the driving force that maintains the HSC fate in the IAHCs.

*Cis* interactions *per se* have been postulated based on mathematical modelling of Notch receptor and ligand interactions. This model of Notch activity fine tunes tissue pattern induction and is further validated as a vital mode of signalling in engineered cell cultures systems ^36–38, 72^. *Cis* inhibition has been shown to prevent receptor activation when ligands are in stochiometric access^73^ and functionally demonstrated in *Drosophila* development of the wing, eye, oogenesis and the notum^74–76^. Disruption of *cis* inhibition by mutating specific extracellular regions of the *Jag* orthologue, *Serrate*, results in the wing vein loss phenotype^77^. Despite the different examples in *Drosophila*, demonstration of similar *cis*-inhibitory mechanisms in vertebrates has been elusive, likely due to the higher complexity of Notch signaling. However, mathematical modelling of the Notch pathway predicts that *cis* interactions are an integral part of Notch signaling also in vertebrate systems and needs further exploration^78^. This study presents the first phenotypic, functional, and visual evidence of such interaction in the vertebrate tissue.

Finally, generating HSC through re-programming of somatic or differentiated cells, or the *in vitro* differentiation of (induced Pluripotent) Embryonic Stem (ES) cells is a goal of regenerative medicine that is still unmet. Indeed, several studies have documented the expression and requirement of Notch signaling molecules for inducing a “definitive” blood precursor population^70, 79–84^. Still, the repopulation capacity of these cells is poor, lineage biased and/or only transient in the recipient^85–91^. Here, we have unraveled an important regulatory mechanism for Notch receptor and ligand interaction through *Rfng*. We speculate that the *cis* (inhibitory) conformation might be a vital mode of regulation that is lacking in *in vitro* systems. Further studies are needed to dissect the modules of Notch signaling in ES cell-based differentiation to blood and try to implement the missing interactions.

## Supporting information

Thambyrajah_et_al_full_manuscript no suppl

## Data availability

Single cell RNA-Seq and nascent RNA-seq data: GEO accession number.

## Acknowledgements

We thank Sarah Bray and David Sprinzak for vital discussions about *cis* interactions. We thank all members of Espinosa and Bigas laboratories for helpful discussions and technical support. We also thank the animal facility, FACS facility, and genomic facility of the PRBB and CRUK Manchester Institute for their technical support. This work was funded by grants from SAF2016-75613-R and PID2019-104695RB-I00 from Agencia Estatal de Investigación (AEI) and SLT002/16/00299 from Department of Health, Generalitat de Cataluña and 2021 SGR 00039 from AGAUR, Generalitat de Catalunya. The work in G.L. laboratory is supported by Blood Cancer UK (19014) and Cancer Research UK Manchester Institute Core Grant (C5759/A27412). RT is a recipient of BP2016 (00021) and BP/MSCA 2018 (00034) fellowship programs from Generalitat de Catalunya/MSCA.

## Author contributions

AB, LE and RT conceptualized the study, designed the experiments, analyzed data and wrote the manuscript. RT, WHN, SG, DG and FM performed experiments and analyzed data. MM-G, KI, YG, ZF and XW analyzed the scRNAseq and nascent RNA-sequencing data. BG and FC supervised the scRNA seq analysis. ME, M-CF, EP, RB, and GL supervised data analysis. RT, AB, GL and LE wrote the manuscript.

## Materials and Methods

### Mouse lines and animal work

The CD1 wild type strain and *Gfi1:tomato* (Thambyrajah et al., 2016) were used in this study. For time matings, *Gfi1:tomatot^omato^*or CD1 WT females were mated to *Gfi1^tomato^* or CD1 WT males. Vaginal plug detection was considered as day 0.5. The resulting embryos were genotyped and used for FACS analysis and sorting, IHC and Index sorting. Animals were kept under pathogen-free conditions, and all procedures were approved by the Animal Care Committee of the Parc de Recerca Biomedica de Barcelona, license number 9309 approved by the Generalitat de Catalunya.

### Genotyping PCR

Small pieces of embryonic tissue or yolk sac were dissected off the embryo and placed in PCR tube containing 100μl of PBS. The tissue pieces were boiled for 8 minutes at 98°C for denaturation and further digested with Proteinase K (50 μg/ml) for 30 minutes at 55°C, and the enzyme deactivated by boiling the samples for a further 10 minutes at 95°C. 1μl of the samples was used as a template for the PCR.

### AGM dissection, single cell suspension

AGMs of E10 - E11.5 embryos were dissected in PBS with 7% fetal calf serum (FBS) and penicillin/streptomycin (100 U/mL). Single cell suspensions were generated by incubating the tissues for 20-30 minutes in 500 ul of 1mg/ml of Collagenase/Dispase (Roche cat# 10269638001) before mechanical dissociation with a syringe and needle. The resulting single cell suspension was used for antibody staining (see Suppl table T4 for list of antibodies). Samples were analysed on a Fortessa instrument or sorted with FACS AriaII or BD Influx (BD Biosciences). FACS plots were generated using FlowJo V10.

### Single-cell RNA sequencing

E11.5 AGMs were prepared for flow sorting as described above. Single cells were sorted into 384-well plates containing lysis buffer and snap frozen. Libraries were prepared using a modified version of the Smart-Seq2 protocol. Briefly, cDNA was prepared using a Mantis platform (Formulatrix) and quantified with quantIT picogreen reagent (Thermo Fisher). Dual indexed sequencing libraries were prepared from 0.1ng cDNA using an Echo525 automation system (Labcyte) in miniaturized reaction volumes. The library pool was quantified by qPCR using a Library Quantification Kit for Illumina sequencing platforms (Kapa Biosystems). Paired-end 75bp sequencing was carried out by clustering 1.5pM of the library pool on a NextSeq 500 sequencer (Illumina).

### scRNASeq data analysis

Raw reads were mapped against the Mus musculus genome (mm10) with STAR aligner tool (v2.7.3) (Dobin et al., 2013). HTSeq framework (v0.9.1) was used for gene expression quantification (Anders et al., 2015). Raw count matrix contained data from 860 sequenced cells and 46,170 genes. Python v3.7.3 and scanpy (v1.4.4) were used for data pre-processing and main downstream analysis (Wolf et al., 2018). Other relevant Python packages included anndata (v0.6.22.post1), umap (v0.3.10), pandas (v0.25.1), scikit-learn (v0.21.3) and statsmodels (v0.13.15). Cells with less than 2,500 expressed genes and less than 50,000 counts were discarded, and no more than 5% mitochondrial gene expression. Genes expressed in at least one cell were kept. No batch correction was performed. After quality control, we obtained an expression matrix including 775 cells and 30,362 genes. Cyclone was used to classify cells into G1, G2 or S cycle phases (Scialdone et al., 2015). Overall gene expression was normalized to 10,000 counts and logarithmically transformed. Then highly variable genes (HVG) were selected with default parameters. A total of 7,925 HVG were identified. Read depth, mitochondrial gene content and cell cycle effects were regressed out. For cells visualization and clustering, PCA was first performed with 50 components on the list of HVG. UMAP was obtained after neighborhood graph was computed (n_neighbors = 5 and 50 PCs). Cells were clustered using the Louvain algorithm with a resolution of 0.5. This analyisis identified 11 clusters. Cell identities annotation to clusters was based on already established marker genes from the literature. Cluster-specific marker genes were found using the scanpy function rank_genes_groups using Wilcoxon rank-sum test method. Only those genes present in at least 25% of cells in either groups (cluster of interest against the rest) were considered.

Benjamini-Hochberg procedure was used to obtain adjusted p-values (Benjamini & Hochberg, 1995). Forced Directed Graph (FDG) was computed after obtaining the diffusion map with default parameters. For the layout, fa2 (v0.3.5) was selected. Cells pseudotime was estimated considering HSC-HE cluster cells as the root.

Data visualization was performed with the ggplot2 (v3.4.1), complexHeatmap (v2.14.0) and EnhancedVolcano (v1.16.0) R packages (v4.2.1) (Gu et al., 2016; R Core Team, 2021). Differential expression analysis (DEA) was performed for cells simultaneously expressing Hey1 and Hey2 genes from GFI1+ HE population (subset of 281 cells). Genes were considered expressed if their number of normalized counts was higher than 0. DEA was independently conducted per time point (E10.5 and E11.5). Same approach was applied as for the cluster-specific markers identification. Those genes with adjusted p-value (FDR) < 0.05 and absolute log2 FC > 2 were identified as differentially expressed genes (DEGs).

### Immuno- histo-chemistry

E10.5 embryos were fixed in 2% Paraformaldehyde (Thermo Fisher) for 12 minutes, before they were soaked in 30% sucrose overnight and mounted in OCT compound. 10-12 µm sections were prepared using a cryostat. The sections were streptavidin/biotin blocked if a biotin antibody was used followed by serum blocking (PBS with 10% FCS, 0.05% Tween20) for 1 hour before the sections were incubated with primary antibodies at 4°C overnight in blocking buffer. Sections were washed three times in PBST for 15 minutes each and then incubated with fluorochrome-conjugated secondary antibody at room temperature for 1 hour. Sections were further washed three times in PBST and mounted using Prolong Gold anti-fade medium with DAPI (Life Technologies). Images were taken using a SPE (Leica) and processed using Imaris v4.8.

### Intracellular FACS staining

RFNG staining was performed on E11.5 AGM cell lysates. Single cell suspension was stained for the cell surface markers before the intracellular (rabbit anti-mouse) RFNG and a secondary antibody (donkey anti rabbit Alexa 488) was performed in FIX & PERM™ Cell Permeabilization Kit from Thermo Fisher (cat# GAS003). The samples were run on a Fortessa Instrument from BD Bioscience and analyzed with FlowJO V10. Statistical significance was determined with GraphPad prism 8.

### Proximity Ligation assay (PLA)

E10.5 and E11.5 CD1 or *Gfi1:tomato* trunks were cut into 150mm thick sections, fixed in 2% Paraformaldehyde (Thermo Fisher) for 12 minutes, and permeabilized for 10 minutes in −20C 100% Methanol. Proximity Ligation assay was performed according to manufacturer’s protocol (Sigma, cat# DUO96000). Briefly, after washing with PBS, the sections were blocked for 1 hour with blocking solution and then incubated with the indicated primary antibody pairs as multiplex with N1/DLL4 (Sigma, Duolink DUO92008), N1ex/JAG1ex (sigma Duolink DUO92013), N1int/JAG1int (sigma, Duolink DUO92014) and (rat anti-mouse) CD31 or (rat anti-mouse) cKIT (please see antibody list in Suppl Table T1). The thick sections were embedded in 80% glycerol in glass bottomed petri dishes (ibidi cat# 81156) and imaged with a SPE (Leica) instrument. Images were taken as z-stacks and rendered to a 3D representation using Imaris v4.8. PLA signals from cell images were manually counted and plotted using Prism (GraphPad prism 8).

### Nascent RNA capture assay

E11.5 *Gfi1:tomato* AGM cell suspension were stained for CD31 and cKIT. Endo (CD31+cKIT-GFI1-) or GFI1+IAHC (CD31+cKIT+GFI1+) were FACS purified and treated with 10μM of compound E (gamma-Secretase Inhibitor XXI,, Merk cat # 565790) overnight. The following morning, the two samples were washed, separated as 300-400 cells/sample and incubated for 4 hours in methylcellulose with the addition 0.3mM of EU (ethynyl Uridine, Thermo Fisher, cat #C10365) with 1ul DMSO, 10μM of compound E or 4μg/ml of recombinant Fc-JAG1 (R&D Systems, Cat #599-JG). After the EU labelling, cells were harvested and processed for purification of EU labeled RNA. Nascent RNA was purified from total nuclear RNA samples using the Click-iT Nascent RNA Capture Kit (Thermo Fisher, cat #C10365) according to the manufacturer’s instructions.

### Nascent RNA capture-sample processing

Nascent RNA was purified from total nuclear RNA samples using the Click-iT Nascent RNA Capture Kit (Thermo Fisher, cat #C10365) according to the manufacturer’s instructions. In brief, biotin-azide was attached to ethylene-groups of the EU-labeled RNA using click-it chemistry and the pulled down of EU-RNA captured on the beads was immediately pre-amplified for 11 cycles (SuperScript™ IV Single Cell/Low Input cDNA PreAmp Kit, cThermo Fisher cat # 11752048) in accordance to manufacturer’s instructions. Finally, double-stranded cDNA was purified using Agencourt AMPure XP beads (Beckman Coulter, cat #A63882). Validation PCRs were performed using the primers listed in Supplemental Table T1.

### Nascent RNASeq data analysis

Libraries were prepared at the Genomics Unit of PRBB (Barcelona, Spain) using Clontech SMARTer kit for low input material and cDNA was sequenced using Illumina NextSeq 2000 platform (50bp single end reads). Samples sequencing depth ranged between 46M and 53M reads (average 49M reads) per sample.

Quality control was performed on raw data with FASTQC tool (v0.11.9). Raw reads were trimmed to remove Clontech SMARTer IIA oligo (AAGCAGTGGTATCAACGCAGAGTAC) 5’ presence with cutadapt (v4.2) (Martin, 2011). Default parameters were used except for a maximum 5% error rate and no indels allowed. Trimmed reads were aligned to reference genome with STAR aligner tool (v2.7.8). STAR was executed with default parameters except for (i) the number of allowed mismatches was set to 1 and (ii) short reads consideration was relaxed to 50% of read length. Required genome index was built with corresponding GRCm38 gtf and fasta files retrieved from Ensembl (http://ftp.ensembl.org/pub/release-102/). Obtained BAM files with uniquely mapped reads were considered for further analysis. Raw gene expression was quantified using featureCounts tool from subRead software (v2.0.1) with gene as feature (Liao et al., 2014). Obtained raw counts matrix was imported into R Statistical Software environment (v4.2.1) for downstream analysis. Raw expression matrix included 55,487 genes per 7 samples which were distributed in 4 different conditions: 2xFc-JAG1, 2xCompE and 3xWashed-out. Prior to statistical analysis, those genes with less than 10 raw counts in at least two replicates from the same condition were removed. After pre-filtering, 10,311 genes were available for testing. For visualization purposes, counts were normalized by variance-stabilizing transformation method using local fit Type as implemented in DESeq2 R package (Love et al., 2014) (v1.38.3). To conduct PCA, normalized expression matrix was corrected for corresponding embryo with the function removeBatchEffect from limma R package (Ritchie et al., 2015) (v3.54.2). Differential expression analysis (DEA) was conducted with DESeq2. Fitted statistical model included embryo and sample condition as covariates. Pairwise comparisons for condition levels were tested considering the washed-out as the reference. DEGs were called with adjusted p-values (FDR) < 0.05 and expressed in at least two replicates in any of the two conditions being compared (minimum of 10 raw counts per sample).

### Functional analysis

Overrepresentation analysis was applied over lists of selected genes from scRNA-seq data or RNA-seq data analysis. The Gene Ontology (Biological Process ontology, GO BP terms) and KEGG PATHWAY databases for Mus Musculus (Ashburner et al., 2000; Kanehisa & Goto, 2000) were interrogated by means of clusterProfiler R package (v4.6.2) (Wu et al., 2021). Corresponding Entrez identifiers were used. Benjamini-Hochberg procedure was used to obtain adjusted p-values. Overrepresented GO BP terms (adjusted p-value < 0.05) were simplified using the simplify function from clusterProfiler with default parameters. A simplified list of terms was plotted on the semantic space obtained from REVIGO (Supek et al., 2011).

### *Ex vivo* culture of AGMs with scramble or RFNG FANA-ASO

AGMs of E10.5 embryos were dissected in PBS with 7% fetal calf serum (FCS) and penicillin/streptomycin (100 U/mL) and cultured as explants for 2 days in medium consisting of Stemspan (Stem Cell Technologies, cat # 09600), 20% fetal calf serum, L-glutamine (4 mM), penicillin/streptomycin (50 units/ml), mercaptoethanol (0.1 mM), IL-3 (100 ng/ml), SCF (100 ng/ml) and Flt3L (100 ng/ml). All growth factors were purchased from Peprotech. Tissues were maintained in 5% CO2 at 37°C. To knock down RFNG, we purchased custom designed ASO-FANA oligos targeting the mRNA of RFNG (Aum Biotech, USA). We used either 10nM of control/scramble or RFNG FANA-ASO during the culture period.

### Quantification and statistical analysis

Statistical parameters, including number of events quantified, standard deviation, and statistical significance, are reported in the figures and in the figure legends. Statistical analysis has been performed using GraphPad Prism 8 software (GraphPad), and P < 0.05 is considered significant. Two-sided Student’s t-test was used to compare differences between two groups.

## References

1. Boisset, J.C., van Cappellen, W., Andrieu-Soler, C., Galjart, N., Dzierzak, E., and Robin, C. (2010). In vivo imaging of haematopoietic cells emerging from the mouse aortic endothelium. Nature 464, 116–120. 10.1038/nature08764.

2. de Bruijn, M.F., Ma, X., Robin, C., Ottersbach, K., Sanchez, M.J., and Dzierzak, E. (2002). Hematopoietic stem cells localize to the endothelial cell layer in the midgestation mouse aorta. Immunity 16, 673–683.

3. Taoudi, S., and Medvinsky, A. (2007). Functional identification of the hematopoietic stem cell niche in the ventral domain of the embryonic dorsal aorta. Proc Natl Acad Sci U S A 104, 9399–9403. 10.1073/pnas.0700984104.

4. Mascarenhas, M.I., Parker, A., Dzierzak, E., and Ottersbach, K. (2009). Identification of novel regulators of hematopoietic stem cell development through refinement of stem cell localization and expression profiling. Blood 114, 4645–4653. 10.1182/blood-2009-06-230037.

5. Yokomizo, T., and Dzierzak, E. (2010). Three-dimensional cartography of hematopoietic clusters in the vasculature of whole mouse embryos. Development 137, 3651–3661. 10.1242/dev.051094.

6. Kumaravelu, P., Hook, L., Morrison, A.M., Ure, J., Zhao, S., Zuyev, S., Ansell, J., and Medvinsky, A. (2002). Quantitative developmental anatomy of definitive haematopoietic stem cells/long-term repopulating units (HSC/RUs): role of the aorta-gonad-mesonephros (AGM) region and the yolk sac in colonisation of the mouse embryonic liver. Development 129, 4891–4899.

7. Boisset, J.C., Clapes, T., Klaus, A., Papazian, N., Onderwater, J., Mommaas-Kienhuis, M., Cupedo, T., and Robin, C. (2015). Progressive maturation toward hematopoietic stem cells in the mouse embryo aorta. Blood 125, 465–469. 10.1182/blood-2014-07-588954.

8. Patel, S.H., Christodoulou, C., Weinreb, C., Yu, Q., da Rocha, E.L., Pepe-Mooney, B.J., Bowling, S., Li, L., Osorio, F.G., Daley, G.Q., et al. (2022). Lifelong multilineage contribution by embryonic-born blood progenitors. Nature 606, 747–753. 10.1038/s41586-022-04804-z.

9. Yokomizo, T., Ideue, T., Morino-Koga, S., Tham, C.Y., Sato, T., Takeda, N., Kubota, Y., Kurokawa, M., Komatsu, N., Ogawa, M., et al. (2022). Independent origins of fetal liver haematopoietic stem and progenitor cells. Nature 609, 779–784. 10.1038/s41586-022-05203-0.

10. Ganuza, M., Hall, T., Finkelstein, D., Chabot, A., Kang, G., and McKinney-Freeman, S. (2017). Lifelong haematopoiesis is established by hundreds of precursors throughout mammalian ontogeny. Nat Cell Biol 19, 1153–1163. 10.1038/ncb3607.

11. Thambyrajah, R., Mazan, M., Patel, R., Moignard, V., Stefanska, M., Marinopoulou, E., Li, Y., Lancrin, C., Clapes, T., Moroy, T., et al. (2016). GFI1 proteins orchestrate the emergence of haematopoietic stem cells through recruitment of LSD1. Nat Cell Biol 18, 21–32. 10.1038/ncb3276ncb3276 [pii].

12. Kauts, M.L., Rodriguez-Seoane, C., Kaimakis, P., Mendes, S.C., Cortes-Lavaud, X., Hill, U., and Dzierzak, E. (2018). In Vitro Differentiation of Gata2 and Ly6a Reporter Embryonic Stem Cells Corresponds to In Vivo Waves of Hematopoietic Cell Generation. Stem Cell Reports 10, 151–165. S2213-6711(17)30526-X [pii] 10.1016/j.stemcr.2017.11.018.

13. Nottingham, W.T., Jarratt, A., Burgess, M., Speck, C.L., Cheng, J.F., Prabhakar, S., Rubin, E.M., Li, P.S., Sloane-Stanley, J., Kong, A.S.J., et al. (2007). Runx1-mediated hematopoietic stem-cell emergence is controlled by a Gata/Ets/SCL-regulated enhancer. Blood 110, 4188–4197. blood-2007-07-100883 [pii] 10.1182/blood-2007-07-100883.

14. Eich, C., Arlt, J., Vink, C.S., Solaimani Kartalaei, P., Kaimakis, P., Mariani, S.A., van der Linden, R., van Cappellen, W.A., and Dzierzak, E. (2018). In vivo single cell analysis reveals Gata2 dynamics in cells transitioning to hematopoietic fate. J Exp Med 215, 233–248. 10.1084/jem.20170807jem.20170807 [pii].

15. Chen, M.J., Yokomizo, T., Zeigler, B.M., Dzierzak, E., and Speck, N.A. (2009). Runx1 is required for the endothelial to haematopoietic cell transition but not thereafter. Nature 457, 887–891. 10.1038/nature07619.

16. Vink, C.S., Calero-Nieto, F.J., Wang, X., Maglitto, A., Mariani, S.A., Jawaid, W., Göttgens, B., and Dzierzak, E. (2020). Iterative Single-Cell Analyses Define the Transcriptome of the First Functional Hematopoietic Stem Cells. Cell Rep 31. 10.1016/j.celrep.2020.107627.

17. Porcheri, C., Golan, O., Calero-Nieto, F.J., Thambyrajah, R., Ruiz-Herguido, C., Wang, X., Catto, F., Guillen, Y., Sinha, R., Gonzalez, J., et al. (2020). Notch ligand Dll4 impairs cell recruitment to aortic clusters and limits blood stem cell generation. EMBO J 39, e104270. 10.15252/embj.2019104270.

18. Fadlullah, M.Z., Neo, W.H., Lie, A.L.M., Thambyrajah, R., Patel, R., Mevel, R., Aksoy, I., Do Khoa, N., Savatier, P., Fontenille, L., et al. (2021). Murine AGM single-cell profiling identifies a continuum of hemogenic endothelium differentiation marked by ACE. Blood. blood.2020007885 [pii] 10.1182/blood.2020007885 S0006-4971(21)01595-0 [pii].

19. Lancrin, C., Mazan, M., Stefanska, M., Patel, R., Lichtinger, M., Costa, G., Vargel, O., Wilson, N.K., Moroy, T., Bonifer, C., et al. (2012). GFI1 and GFI1B control the loss of endothelial identity of hemogenic endothelium during hematopoietic commitment. Blood 120, 314–322. 10.1182/blood-2011-10-386094.

20. Boisset, J.C., van Cappellen, W., Andrieu-Soler, C., Galjart, N., Dzierzak, E., and Robin, C. (2010). In vivo imaging of haematopoietic cells emerging from the mouse aortic endothelium. Nature 464, 116–120. 10.1038/nature08764.

21. Bertrand, J.Y., Chi, N.C., Santoso, B., Teng, S., Stainier, D.Y., and Traver, D. (2010). Haematopoietic stem cells derive directly from aortic endothelium during development. Nature 464, 108–111. 10.1038/nature08738.

22. Kissa, K., and Herbomel, P. (2010). Blood stem cells emerge from aortic endothelium by a novel type of cell transition. Nature 464, 112–115. 10.1038/nature08761.

23. Medvinsky, A., Rybtsov, S., and Taoudi, S. (2011). Embryonic origin of the adult hematopoietic system: advances and questions. Development 138, 1017–1031. 10.1242/dev.040998.

24. Rybtsov, S., Sobiesiak, M., Taoudi, S., Souilhol, C., Senserrich, J., Liakhovitskaia, A., Ivanovs, A., Frampton, J., Zhao, S., and Medvinsky, A. (2011). Hierarchical organization and early hematopoietic specification of the developing HSC lineage in the AGM region. J Exp Med 208, 1305–1315. 10.1084/jem.20102419.

25. Taoudi, S., Gonneau, C., Moore, K., Sheridan, J.M., Blackburn, C.C., Taylor, E., and Medvinsky, A. (2008). Extensive hematopoietic stem cell generation in the AGM region via maturation of VE-cadherin+CD45+ pre-definitive HSCs. Cell Stem Cell 3, 99–108. 10.1016/j.stem.2008.06.004.

26. Zhou, F., Li, X., Wang, W., Zhu, P., Zhou, J., He, W., Ding, M., Xiong, F., Zheng, X., Li, Z., et al. (2016). Tracing haematopoietic stem cell formation at single-cell resolution. Nature 533, 487–492. 10.1038/nature17997.

27. Hadland, B.K., Varnum-Finney, B., Nourigat-Mckay, C., Flowers, D., and Bernstein, I.D. (2018). Clonal analysis of embryonic hematopoietic stem cell precursors using single cell index sorting combined with endothelial cell niche co-culture. Journal of Visualized Experiments 2018. 10.3791/56973.

28. Baron, C.S., Kester, L., Klaus, A., Boisset, J.C., Thambyrajah, R., Yvernogeau, L., Kouskoff, V., Lacaud, G., van Oudenaarden, A., and Robin, C. (2018). Single-cell transcriptomics reveal the dynamic of haematopoietic stem cell production in the aorta. Nat Commun 9, 2517. 10.1038/s41467-018-04893-3 10.1038/s41467-018-04893-3 [pii].

29. Zhu, Q., Gao, P., Tober, J., Bennett, L., Chen, C., Uzun, Y., Li, Y., Howell, E.D., Mumau, M., Yu, W., et al. (2020). Developmental trajectory of prehematopoietic stem cell formation from endothelium. Blood 136, 845–856. 10.1182/blood.2020004801.

30. Zhou, F., Li, X., Wang, W., Zhu, P., Zhou, J., He, W., Ding, M., Xiong, F., Zheng, X., Li, Z., et al. (2016). Tracing haematopoietic stem cell formation at single-cell resolution. Nature 533, 487–492. 10.1038/nature17997.

31. Artavanis-Tsakonas, S., Rand, M.D., and Lake, R.J. (1999). Notch signaling: cell fate control and signal integration in development. Science (1979) 284, 770–776. 10.1126/science.284.5415.770.

32. Gazave, E., Lapébie, P., Richards, G.S., Brunet, F., Ereskovsky, A. V., Degnan, B.M., Borchiellini, C., Vervoort, M., and Renard, E. (2009). Origin and evolution of the Notch signalling pathway: An overview from eukaryotic genomes. BMC Evol Biol 9. 10.1186/1471-2148-9-249.

33. Kageyama, R., Ohtsuka, T., and Kobayashi, T. (2007). The Hes gene family: Repressors and oscillators that orchestrate embryogenesis. Development 134, 1243–1251. 10.1242/dev.000786.

34. Bray, S.J. (2016). Notch signalling in context. Nat Rev Mol Cell Biol 17, 722–735. 10.1038/nrm.2016.94.

35. Gozlan, O., and Sprinzak, D. (2023). Notch signaling in development and homeostasis. Development 150, dev201138. 10.1242/dev.201138.

36. Nandagopal, N., Santat, L.A., and Elowitz, M.B. (2019). Cis-activation in the Notch signaling pathway. Elife 8. 10.7554/eLife.37880.

37. Sprinzak, D., Lakhanpal, A., Lebon, L., Santat, L.A., Fontes, M.E., Anderson, G.A., Garcia-Ojalvo, J., and Elowitz, M.B. (2010). Cis-interactions between Notch and Delta generate mutually exclusive signalling states. Nature 465, 86–90. 10.1038/nature08959.

38. Del Álamo, D., Rouault, H., and Schweisguth, F. (2011). Mechanism and significance of cis-inhibition in notch signalling. Current Biology 21. 10.1016/j.cub.2010.10.034.

39. Xu, X., Seymour, P.A., Sneppen, K., Trusina, A., Egeskov-Madsen, A. la R., Jørgensen, M.C., Jensen, M.H., and Serup, P. (2023). Jag1-Notch cis-interaction determines cell fate segregation in pancreatic development. Nat Commun 14, 348. 10.1038/s41467-023-35963-w.

40. Takeuchi, H., and Haltiwanger, R.S. (2014). Significance of glycosylation in Notch signaling. Biochem Biophys Res Commun 453, 235–242. 10.1016/j.bbrc.2014.05.115.

41. LeBon, L., Lee, T. V., Sprinzak, D., Jafar-Nejad, H., and Elowitz, M.B. (2014). Fringe proteins modulate Notch-ligand cis and trans interactions to specify signaling states. Elife 3, e02950. 10.7554/eLife.02950.

42. Thambyrajah, R., and Bigas, A. (2022). Notch Signaling in HSC Emergence: When, Why and How. Cells 11, 358. 10.3390/cells11030358.

43. Hadland, B.K., Huppert, S.S., Kanungo, J., Xue, Y., Jiang, R., Gridley, T., Conlon, R.A., Cheng, A.M., Kopan, R., and Longmore, G.D. (2004). A requirement for Notch1 distinguishes 2 phases of definitive hematopoiesis during development. Blood 104, 3097–3105. 10.1182/blood-2004-03-1224.

44. Kumano, K., Chiba, S., Kunisato, A., Sata, M., Saito, T., Nakagami-Yamaguchi, E., Yamaguchi, T., Masuda, S., Shimizu, K., Takahashi, T., et al. (2003). Notch1 but not Notch2 is essential for generating hematopoietic stem cells from endothelial cells. Immunity 18, 699–711. 10.1016/S1074-7613(03)00117-1.

45. Gama-Norton, L., Ferrando, E., Ruiz-Herguido, C., Liu, Z., Guiu, J., Islam, A.B.M.M.K., Lee, S.U., Yan, M., Guidos, C.J., López-Bigas, N., et al. (2015). Notch signal strength controls cell fate in the haemogenic endothelium. Nat Commun 6. 10.1038/ncomms9510.

46. Porcheri, C., Golan, O., Calero-Nieto, F.J., Thambyrajah, R., Ruiz-Herguido, C., Wang, X., Catto, F., Guillen, Y., Sinha, R., Gonzalez, J., et al. (2020). Notch ligand Dll4 impairs cell recruitment to aortic clusters and limits blood stem cell generation. EMBO J 39, e104270. 10.15252/embj.2019104270.

47. Robert-Moreno, À., Guiu, J., Ruiz-Herguido, C., López, M.E., Inglés-Esteve, J., Riera, L., Tipping, A., Enver, T., Dzierzak, E., Gridley, T., et al. (2008). Impaired embryonic haematopoiesis yet normal arterial development in the absence of the Notch ligand Jagged1. EMBO Journal 27, 1886–1895. 10.1038/emboj.2008.113.

48. Laruy, B., Garcia-Gonzalez, I., Casquero-Garcia, V., and Benedito, R. (2020). Endothelial-to-hematopoietic transition is induced by Notch glycosylation and upregulation of Mycn. bioRxiv, 2020.09.13.295238. 10.1101/2020.09.13.295238.

49. Souilhol, C., Lendinez, J.G., Rybtsov, S., Murphy, F., Wilson, H., Hills, D., Batsivari, A., Binagui-Casas, A., McGarvey, A.C., MacDonald, H.R., et al. (2016). Developing HSCs become Notch independent by the end of maturation in the AGM region. Blood 128, 1567–1577. 10.1182/blood-2016-03-708164.

50. Gama-Norton, L., Ferrando, E., Ruiz-Herguido, C., Liu, Z., Guiu, J., Islam, A.B.M.M.K., Lee, S.U., Yan, M., Guidos, C.J., López-Bigas, N., et al. (2015). Notch signal strength controls cell fate in the haemogenic endothelium. Nat Commun 6. 10.1038/ncomms9510.

51. Zhou, F., Li, X., Wang, W., Zhu, P., Zhou, J., He, W., Ding, M., Xiong, F., Zheng, X., Li, Z., et al. (2016). Tracing haematopoietic stem cell formation at single-cell resolution. Nature 533, 487–492. 10.1038/nature17997.

52. Kieusseian, A., de la Grange, P.B., Burlen-Defranoux, O., Godin, I., and Cumano, A. (2012). Immature hematopoietic stem cells undergo maturation in the fetal liver. Development (Cambridge) 139. 10.1242/dev.079210.

53. Cumano, A., Ferraz, J.C., Klaine, M., Di Santo, J.P., and Godin, I. (2001). Intraembryonic, but not yolk sac hematopoietic precursors, isolated before circulation, provide long-term multilineage reconstitution. Immunity 15, 477–485.

54. Kumano, K., Chiba, S., Kunisato, A., Sata, M., Saito, T., Nakagami-Yamaguchi, E., Yamaguchi, T., Masuda, S., Shimizu, K., Takahashi, T., et al. (2003). Notch1 but not Notch2 is essential for generating hematopoietic stem cells from endothelial cells. Immunity 18, 699–711. 10.1016/S1074-7613(03)00117-1.

55. Robert-Moreno, À., Espinosa, L., de la Pompa, J.L., and Bigas, A. (2005). RBPjκ-dependent Notch function regulates Gata2 and is essential for the formation of intra-embryonic hematopoietic cells. Development 132, 1117–1126. 10.1242/dev.01660.

56. Burns, C.E., Traver, D., Mayhall, E., Shepard, J.L., and Zon, L.I. (2005). Hematopoietic stem cell fate is established by the Notch-Runx pathway. Genes Dev 19, 2331–2342. 10.1101/gad.1337005.

57. Guiu, J., Shimizu, R., D’Altri, T., Fraser, S.T., Hatakeyama, J., Bresnick, E.H., Kageyama, R., Dzierzak, E., Yamamoto, M., Espinosa, L., et al. (2013). Hes repressors are essential regulators of hematopoietic stem cell development downstream of notch signaling. Journal of Experimental Medicine 210, 71–84. 10.1084/jem.20120993.

58. Duarte, A., Hirashima, M., Benedito, R., Trindade, A., Diniz, P., Bekman, E., Costa, L., Henrique, D., and Rossant, J. (2004). Dosage-sensitive requirement for mouse Dll4 in artery development. Genes Dev 18, 2474–2478. 10.1101/gad.1239004.

59. Krebs, L.T., Shutter, J.R., Tanigaki, K., Honjo, T., Stark, K.L., and Gridley, T. (2004). Haploinsufficient lethality and formation of arteriovenous malformations in Notch pathway mutants. Genes Dev 18, 2469–2473. 10.1101/gad.1239204.

60. Gale, N.W., Dominguez, M.G., Noguera, I., Pan, L., Hughes, V., Valenzuela, D.M., Murphy, A.J., Adams, N.C., Lin, H.C., Holash, J., et al. (2004). Haploinsufficiency of delta-like 4 ligand results in embryonic lethality due to major defects in arterial and vascular development. Proc Natl Acad Sci U S A 101, 15949–15954. 10.1073/pnas.0407290101.

61. Krebs, L.T., Xue, Y., Norton, C.R., Shutter, J.R., Maguire, M., Sundberg, J.P., Gallahan, D., Closson, V., Kitajewski, J., Callahan, R., et al. (2000). Notch signaling is essential for vascular morphogenesis in mice. Genes Dev 14, 1343–1352. 10.1101/gad.14.11.1343.

62. Porcheri, C., Golan, O., Calero-Nieto, F.J., Thambyrajah, R., Ruiz-Herguido, C., Wang, X., Catto, F., Guillen, Y., Sinha, R., Gonzalez, J., et al. (2020). Notch ligand Dll4 impairs cell recruitment to aortic clusters and limits blood stem cell generation. EMBO J 39, e104270. 10.15252/embj.2019104270.

63. Lizama, C.O., Hawkins, J.S., Schmitt, C.E., Bos, F.L., Zape, J.P., Cautivo, K.M., Borges Pinto, H., Rhyner, A.M., Yu, H., Donohoe, M.E., et al. (2015). Repression of arterial genes in hemogenic endothelium is sufficient for haematopoietic fate acquisition. Nat Commun 6. 10.1038/ncomms8739.

64. Richard, C., Drevon, C., Canto, P.Y., Villain, G., Bollérot, K., Lempereur, A., Teillet, M.A., Vincent, C., Rosselló Castillo, C., Torres, M., et al. (2013). Endothelio-Mesenchymal Interaction Controls runx1 Expression and Modulates the notch Pathway to Initiate Aortic Hematopoiesis. Dev Cell 24, 600–611. 10.1016/j.devcel.2013.02.011.

65. Robert-Moreno, À., Guiu, J., Ruiz-Herguido, C., López, M.E., Inglés-Esteve, J., Riera, L., Tipping, A., Enver, T., Dzierzak, E., Gridley, T., et al. (2008). Impaired embryonic haematopoiesis yet normal arterial development in the absence of the Notch ligand Jagged1. EMBO Journal 27, 1886–1895. 10.1038/emboj.2008.113.

66. Laruy, B., Garcia-Gonzalez, I., Casquero-Garcia, V., and Benedito, R. (2020). Endothelial-to-hematopoietic transition is induced by Notch glycosylation and upregulation of Mycn. bioRxiv, 2020.09.13.295238. 10.1101/2020.09.13.295238.

67. Zhang, P., He, Q., Chen, D., Liu, W., Wang, L., Zhang, C., Ma, D., Li, W., Liu, B., and Liu, F. (2015). G protein-coupled receptor 183 facilitates endothelial-to-hematopoietic transition via Notch1 inhibition. Cell Res 25. 10.1038/cr.2015.109.

68. Souilhol, C., Lendinez, J.G., Rybtsov, S., Murphy, F., Wilson, H., Hills, D., Batsivari, A., Binagui-Casas, A., McGarvey, A.C., MacDonald, H.R., et al. (2016). Developing HSCs become Notch independent by the end of maturation in the AGM region. Blood 128, 1567–1577. 10.1182/blood-2016-03-708164.

69. LeBon, L., Lee, T. V., Sprinzak, D., Jafar-Nejad, H., and Elowitz, M.B. (2014). Fringe proteins modulate Notch-ligand cis and trans interactions to specify signaling states. Elife 3, e02950. 10.7554/eLife.02950.

70. Uenishi, G.I., Jung, H.S., Kumar, A., Park, M.A., Hadland, B.K., McLeod, E., Raymond, M., Moskvin, O., Zimmerman, C.E., Theisen, D.J., et al. (2018). NOTCH signaling specifies arterial-type definitive hemogenic endothelium from human pluripotent stem cells. Nat Commun 9. 10.1038/s41467-018-04134-7.

71. Dignum, T., Varnum-Finney, B., Srivatsan, S.R., Dozono, S., Waltner, O., Heck, A.M., Ishida, T., Nourigat-McKay, C., Jackson, D.L., Rafii, S., et al. (2021). Multipotent progenitors and hematopoietic stem cells arise independently from hemogenic endothelium in the mouse embryo. Cell Rep 36. 10.1016/j.celrep.2021.109675.

72. Sprinzak, D., Lakhanpal, A., LeBon, L., Garcia-Ojalvo, J., and Elowitz, M.B. (2011). Mutual inactivation of Notch receptors and ligands facilitates developmental patterning. PLoS Comput Biol 7. 10.1371/journal.pcbi.1002069.

73. Jacobsen, T.L., Brennan, K., Arias, A.M., and Muskavitch, M.A.T. (1998). Cis-interactions between Delta and Notch modulate neurogenic signalling in Drosophila. Development 125. 10.1242/dev.125.22.4531.

74. De Celis, J.F., and Bray, S. (1997). Feed-back mechanisms affecting Notch activation at the dorsoventral boundary in the Drosophila wing. Development 124, 3241–3251. 10.1242/dev.124.17.3241.

75. Micchelli, C.A., Rulifson, E.J., and Blair, S.S. (1997). The function and regulation of cut expression on the wing margin of Drosophila: Notch, Wingless and a dominant negative role for Delta and Serrate. Development 124, 1485–1495. 10.1242/dev.124.8.1485.

76. Becam, I., Fiuza, U.M., Arias, A.M., and Milán, M. (2010). A Role of Receptor Notch in Ligand cis-Inhibition in Drosophila. Current Biology 20, 554–560. 10.1016/j.cub.2010.01.058.

77. Fleming, R.J., Hori, K., Sen, A., Filloramo, G. V., Langer, J.M., Obar, R.A., Artavanis-Tsakonas, S., and Maharaj-Best, A.C. (2013). An extracellular region of Serrate is essential for ligandinduced cis-inhibition of Notch signaling. Development (Cambridge) 140. 10.1242/dev.087916.

78. Xu, X., Seymour, P.A., Sneppen, K., Trusina, A., Egeskov-Madsen, A. la R., Jørgensen, M.C., Jensen, M.H., and Serup, P. (2023). Jag1-Notch cis-interaction determines cell fate segregation in pancreatic development. Nat Commun 14, 348. 10.1038/s41467-023-35963-w.

79. Ayllon, V., Bueno, C., Ramos-Mejia, V., Navarro-Montero, O., Prieto, C., Real, P.J., Romero, T., Garcia-Leon, M.J., Toribio, M.L., Bigas, A., et al. (2015). The Notch ligand DLL4 specifically marks human hematoendothelial progenitors and regulates their hematopoietic fate. Leukemia 29, 1741–1753. 10.1038/leu.2015.74leu201574 [pii].

80. Slukvin II (2016). Generating human hematopoietic stem cells in vitro -exploring endothelial to hematopoietic transition as a portal for stemness acquisition. FEBS Lett 590, 4126–4143. 10.1002/1873-3468.12283.

81. Jung, H.S., Uenishi, G., Park, M.A., Liu, P., Suknuntha, K., Raymond, M., Choi, Y.J., Thomson, J.A., Ong, I.M., and Slukvin, I.I. (2021). SOX17 integrates HOXA and arterial programs in hemogenic endothelium to drive definitive lympho-myeloid hematopoiesis. Cell Rep 34. 10.1016/j.celrep.2021.108758.

82. Park, M.A., Kumar, A., Jung, H.S., Uenishi, G., Moskvin, O. V., Thomson, J.A., and Slukvin, I.I. (2018). Activation of the Arterial Program Drives Development of Definitive Hemogenic Endothelium with Lymphoid Potential. Cell Rep 23. 10.1016/j.celrep.2018.04.092.

83. Ditadi, A., Sturgeon, C.M., Tober, J., Awong, G., Kennedy, M., Yzaguirre, A.D., Azzola, L., Ng, E.S., Stanley, E.G., French, D.L., et al. (2015). Human definitive haemogenic endothelium and arterial vascular endothelium represent distinct lineages. Nat Cell Biol 17. 10.1038/ncb3161.

84. Lee, J.B., Werbowetski-Ogilvie, T.E., Lee, J.H., McIntyre, B.A.S., Schnerch, A., Hong, S.H., Park, I.H., Daley, G.Q., Bernstein, I.D., and Bhatia, M. (2013). Notch-HES1 signaling axis controls hemato-endothelial fate decisions of human embryonic and induced pluripotent stem cells. Blood 122. 10.1182/blood-2012-12-471649.

85. Wang, L., Menendez, P., Shojaei, F., Li, L., Mazurier, F., Dick, J.E., Cerdan, C., Levac, K., and Bhatia, M. (2005). Generation of hematopoietic repopulating cells from human embryonic stem cells independent of ectopic HOXB4 expression. Journal of Experimental Medicine 201. 10.1084/jem.20041888.

86. Varnum-Finney, B., Brashem-Stein, C., and Bernstein, I.D. (2003). Combined effects of Notch signaling and cytokines induce a multiple log increase in precursors with lymphoid and myeloid reconstituting ability. Blood 101. 10.1182/blood-2002-06-1862.

87. Delaney, C., Varnum-Finney, B., Aoyama, K., Brashem-Stein, C., and Bernstein, I.D. (2005). Dose-dependent effects of the Notch ligand Delta1 on ex vivo differentiation and in vivo marrow repopulating ability of cord blood cells. Blood 106. 10.1182/blood-2005-03-1131.

88. Ohishi, K., Varnum-Finney, B., and Bernstein, I.D. (2002). Delta-1 enhances marrow and thymus repopulating ability of human CD34+CD38-cord blood cells. Journal of Clinical Investigation 110. 10.1172/JCI0216167.

89. Delaney, C., Heimfeld, S., Brashem-Stein, C., Voorhies, H., Manger, R.L., and Bernstein, I.D. (2010). Notch-mediated expansion of human cord blood progenitor cells capable of rapid myeloid reconstitution. Nat Med 16. 10.1038/nm.2080.

90. Suzuki, N., Yamazaki, S., Yamaguchi, T., Okabe, M., Masaki, H., Takaki, S., Otsu, M., and Nakauchi, H. (2013). Generation of engraftable hematopoietic stem cells from induced pluripotent stem cells by way of teratoma formation. Molecular Therapy 21. 10.1038/mt.2013.71.

91. Amabile, G., Welner, R.S., Nombela-Arrieta, C., D’Alise, A.M., Di Ruscio, A., Ebralidze, A.K., Kraytsberg, Y., Ye, M., Kocher, O., Neuberg, D.S., et al. (2013). In vivo generation of transplantable human hematopoietic cells from induced pluripotent stem cells. Blood 121. 10.1182/blood-2012-06-434407.

## References

Dobin, A., Davis, C. A., Schlesinger, F., Drenkow, J., Zaleski, C., Jha, S., Batut, P., Chaisson, M., & Gingeras, T. R. (2013). STAR: ultrafast universal RNA-seq aligner. Bioinformatics (Oxford, England), 29(1), 15–21. https://doi.org/10.1093/bioinformatics/bts635

Liao, Y., Smyth, G. K., & Shi, W. (2014). featureCounts: an efficient general purpose program for assigning sequence reads to genomic features. Bioinformatics (Oxford, England), 30(7), 923–930. https://doi.org/10.1093/bioinformatics/btt656

Love, M. I., Huber, W., & Anders, S. (2014). Moderated estimation of fold change and dispersion for RNA-seq data with DESeq2. Genome Biology, 15(12), 550. https://doi.org/10.1186/s13059-014-0550-8

Martin, M. (2011). Cutadapt removes adapter sequences from high-throughput sequencing reads. EMBnet.Journal; Vol 17, No 1: Next Generation Sequencing Data AnalysisDO *-* 10.14806/Ej.17.1.200. https://journal.embnet.org/index.php/embnetjournal/article/view/200

R Core Team. (2021). R: A Language and Environment for Statistical Computing. R Foundation for Statistical Computing. https://www.r-project.org/

