## Supplementary material for "Radical fringe facilitates NOTCH1 and JAG1 *cis* interactions to sustain Hematopoietic stem cell fate": Thambyrajah_et_al_full_manuscript no suppl

#### Suppl Figure legends

**Suppl Figure S1: Dynamic Notch signaling molecules expression patterns in the AGM** (**A**) Gating for Notch receptors (NOTCH1-4) and ligands (DLL4, JAG1/2) in AGM lysates. The first gate separated endothelial cells (CD31+cKIT-, orange) from IAHC (CD31+CKIT+, blue). The positive population determined in endothelial or IAHC are highlighted in the according color. (**B**) The CD31+cKIT+ IAHC were sub-gated for CD45 expression. The notch receptor and ligand levels within the CD45 positive cells (pink) are highlighted for NOTCH1-4, DLL4 and JAG1/2. (**C**) Bar chart summarizing the expression levels for NOTCH1-4, DLL4 and JAG1/2 within CD31+cKIT- endothelium, CD31+cKIT+ IAHC and CD31+cKIT+CD45+ HSC containing IAHC (n>3 independent experiments with 2-5 AGMs each per sample). Statistical significance was calculated with two-tailed t-tests.

**Suppl Figure S2: GFI1 transgene expression restricts AGM population to HE and HSC containing IAHC** (**A**) Representative Flow Cytometry plots of NOTCH1, DLL4 and JAG1. GFI1 positive cells within CD31cKIT- (HE, green) and CD31+cKIT+ (GFI1+IAHC, purple) were sub gated for NOTCH1, DLL4 and JAG1 in 3132s (E10.25) and 45-47s (E11.5) AGM cell lysates.

**Suppl Figure S3: GFI1+ IAHC retain NOTCH1 and JAG1 co-expression** (**A**) Representative Flow Cytometry plots at 32/33s (E10.5) and 45-48s (E11.5) of CD31+cKIT-GFI1+ (HE, green), CD31+cKIT+GFI1+ (GFI1+IAHC, magenta) and CD31+cKIT+GFI1-(GFI1-IAHC, blue) for JAG1 and DLL4. (**B**) Exemplary Flow Cytometry plots at 32/33s (E10.5) and 45-48s (E11.5) of CD31+cKIT-GFI1+ (HE, green), CD31+cKIT+GFI1+ (GFI1+IAHC, magenta) and CD31+cKIT+GFI1-(GFI1-IAHC, blue) for NOTCH1 and JAG1. (**C**) Representative Flow Cytometry plots at 31/32s (E10.25) and 45-47s (E11.5) of CD31+cKIT+GFI1+ (GFI1+IAHC, magenta) for CD45.

**Suppl Figure S4: Gating and Fluorescence minus one (FMO) control for FACS analysis** Gating strategy to detect CD45, NOTCH1, NOTCH2, JAG1 and DLL4 in endo/HE and IAHC. FMOs were used to set up and verify gating for Flow cytometry analysis.

**Suppl Figure S5: The HSC cluster corresponds to the T2-HSCs** (**A**) UMAP layout colored by developmental stage: E10.5 (orange) and E11.5 (brown). (**B**) UMAP layout representation of inferred cell cycle stage: G1 (yellow), G2/M (green) and S phase (red). (**C**) Heatmap of the top 25 representative genes in each cluster. (**D**) Pseudo-time analysis on all sequenced cells with indication of the cluster identity. (**E**) T1/T2 common genes (98) from Zhou et al, 2016 were plotted for their expression in all clusters with indication of developmental stage and identified clusters (**F**) The gene expression level of *Flt3* across the UMAP representation (**G**) Heatmaps with Index sort levels (left) and gene expression levels (left) for the indicated molecules. Time points (E10.5 orange and E11.5 brown) and identified clusters annotations are included in the top banner.

**Suppl Figure S6: cKIT positive IAHC accumulate NOTCH1-JAG1 in *cis* conformation** (**A**) Immuno-histochemistry for JAG1 (green), DLL4 (red) on GFI1+Tomato (magenta) AGM sections. The localization of DLL4 at cell-cell boundaries are highlighted with a white arrow. Scale bar= 10μm. (**B**) Negative control for Proximity ligation assay. The NOTCH1ex (N1ex) antibody was omitted in this experiment. (**C**) 3D reconstruction from individual z-stacks of the dorsal aorta of E11.5 AGM with proximity ligation assay for NOTCH1ex/JAG1ex (magenta) and NOTCH1/DLL4 (yellow) with CD31 IHC (red) scale bar= 50μm. (**D**) *Tie2: Mfng* overexpressing dorsal aorta assayed with proximity ligation assay for NOTCH1ex/JAG1ex (magenta) and NOTCH1/DLL4 (yellow). Individual z-stacks of 2 different AGM aortas are shown. Scale bar= 10μm. (**E**) Compilation of z-stacks to a 3D representation of a cKIT (red) positive IAHC and DAPI (blue) probed with NOTCH1ex/JAG1ex (magenta) and NOTCH1int/JAG1int (green). Scale bar= 20um. (**F**) Principal component plot of nascent RNA samples with compE treatment (I_C1 and I_C2), after washout (I_W1-I_W3) and stimulation with Fc-JAG1 (I_J1 and I_J3).

**Suppl Figure S7: Radical fringe expression is required for NOTCH1-JAG1 cis conformation and T2-HSC emergence** (**A**) Exemplary Flow cytometry plots for determining CD45+ and CD45- IAHC. (**B**) Representative Flow cytometry plots of scramble and RFNG ASO mediated knock down in AGM explants. RFNG protein levels were assessed after gating on CD31+cKIT+ AGM cells or after further sub gating for CD45.

### Suppl Figure S1

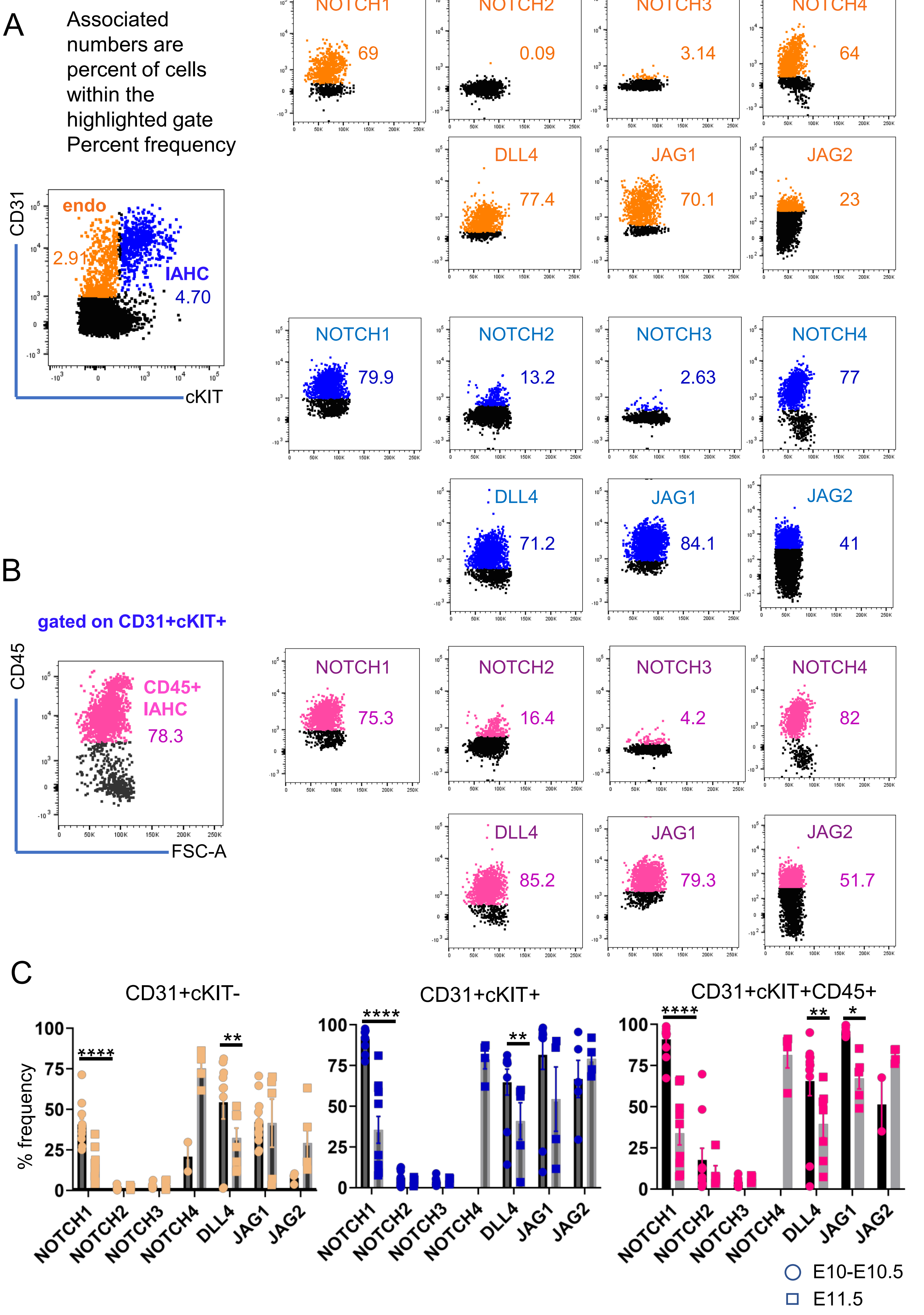

### Suppl Figure S2

A

Associated numbers are percent of cells within the highlighted gate
Percent frequency

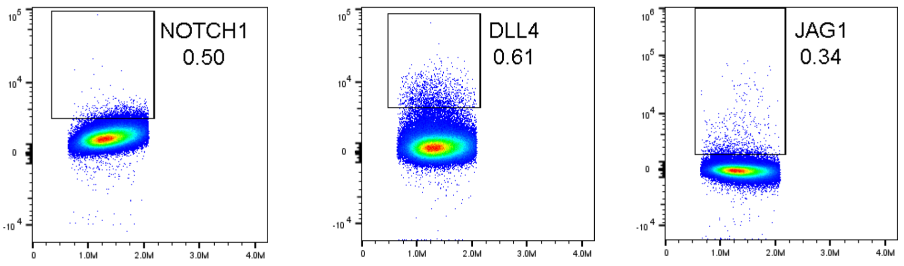


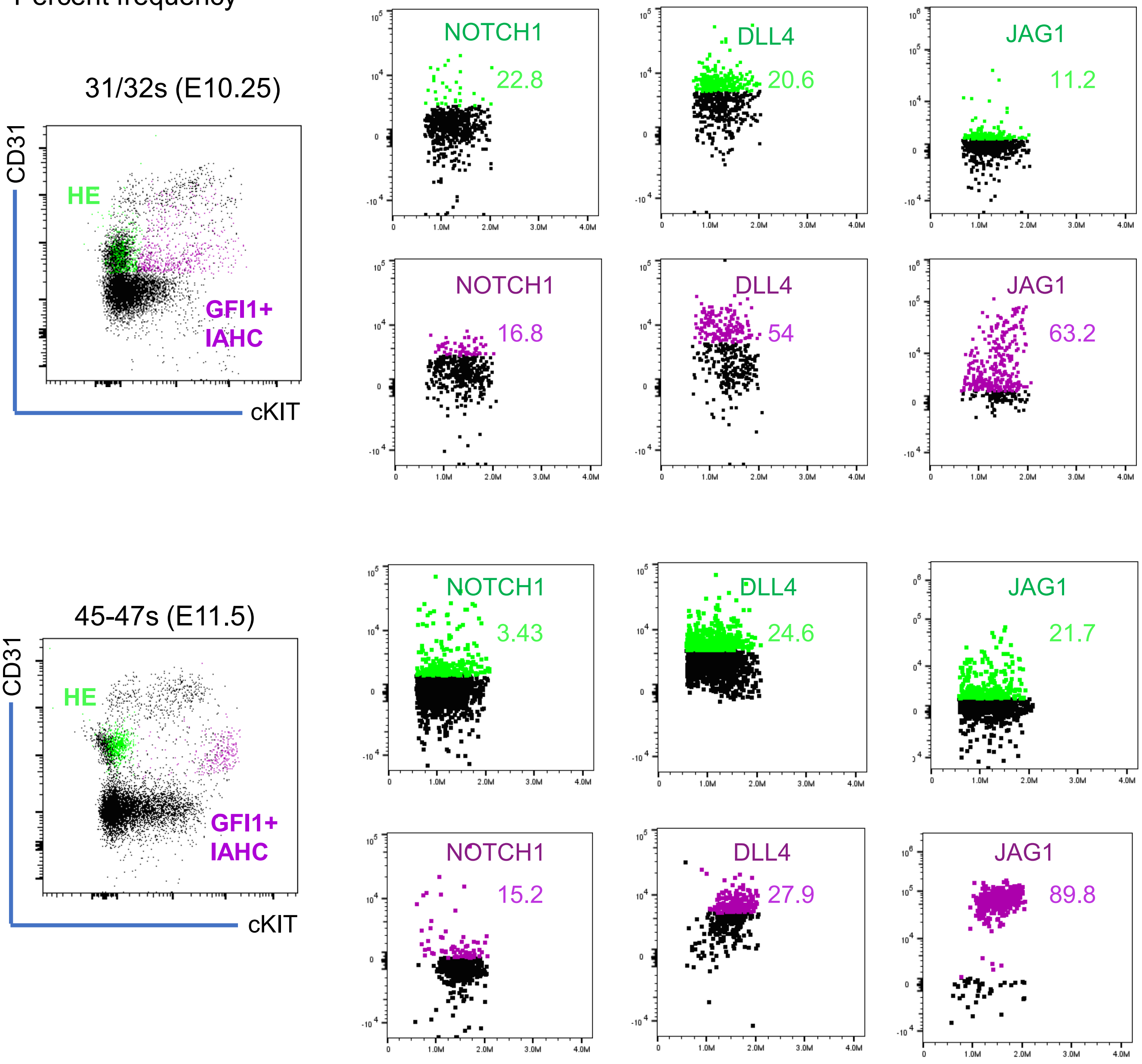

### Suppl Figure S3

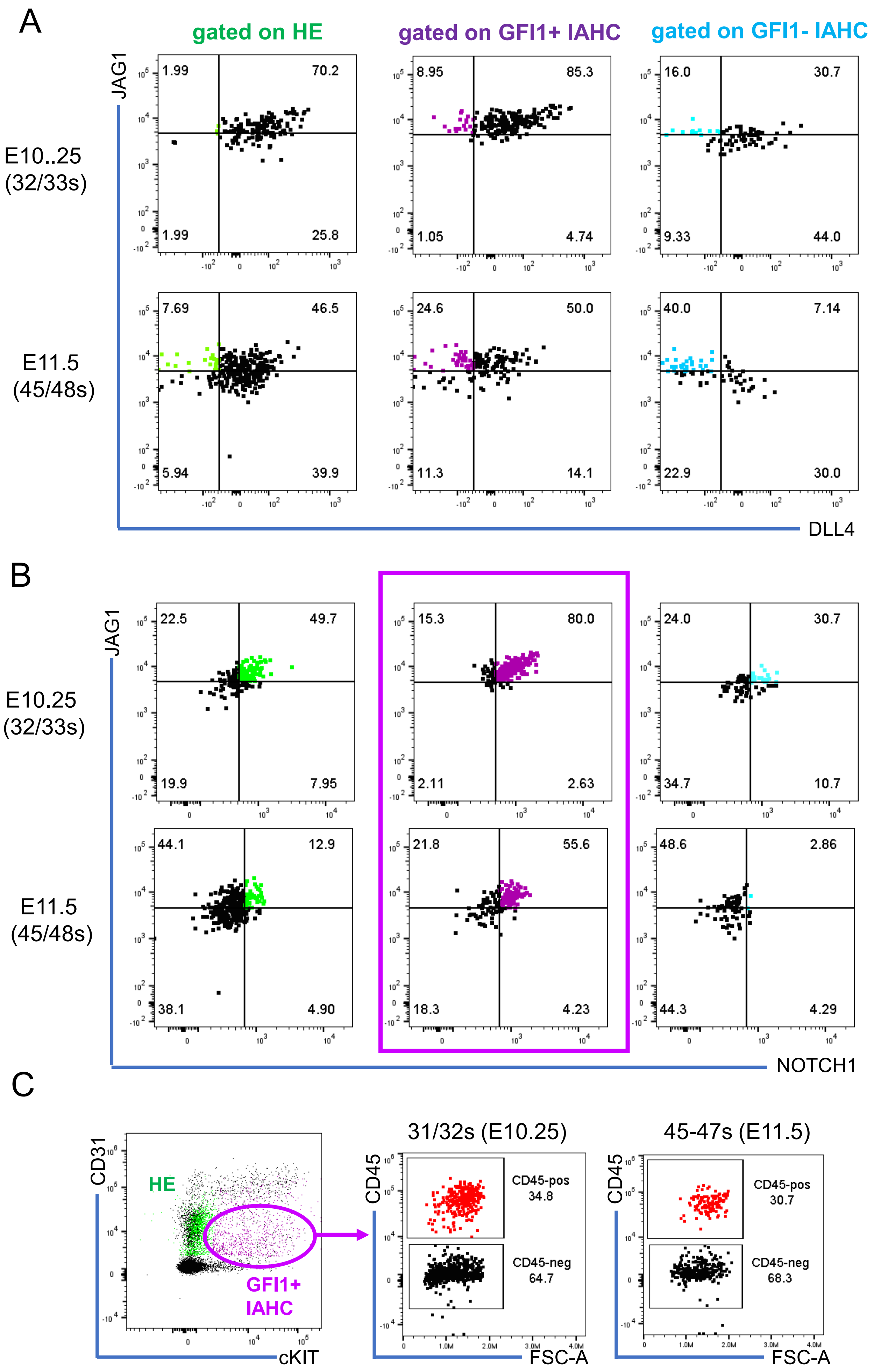

Suppl Figure S4

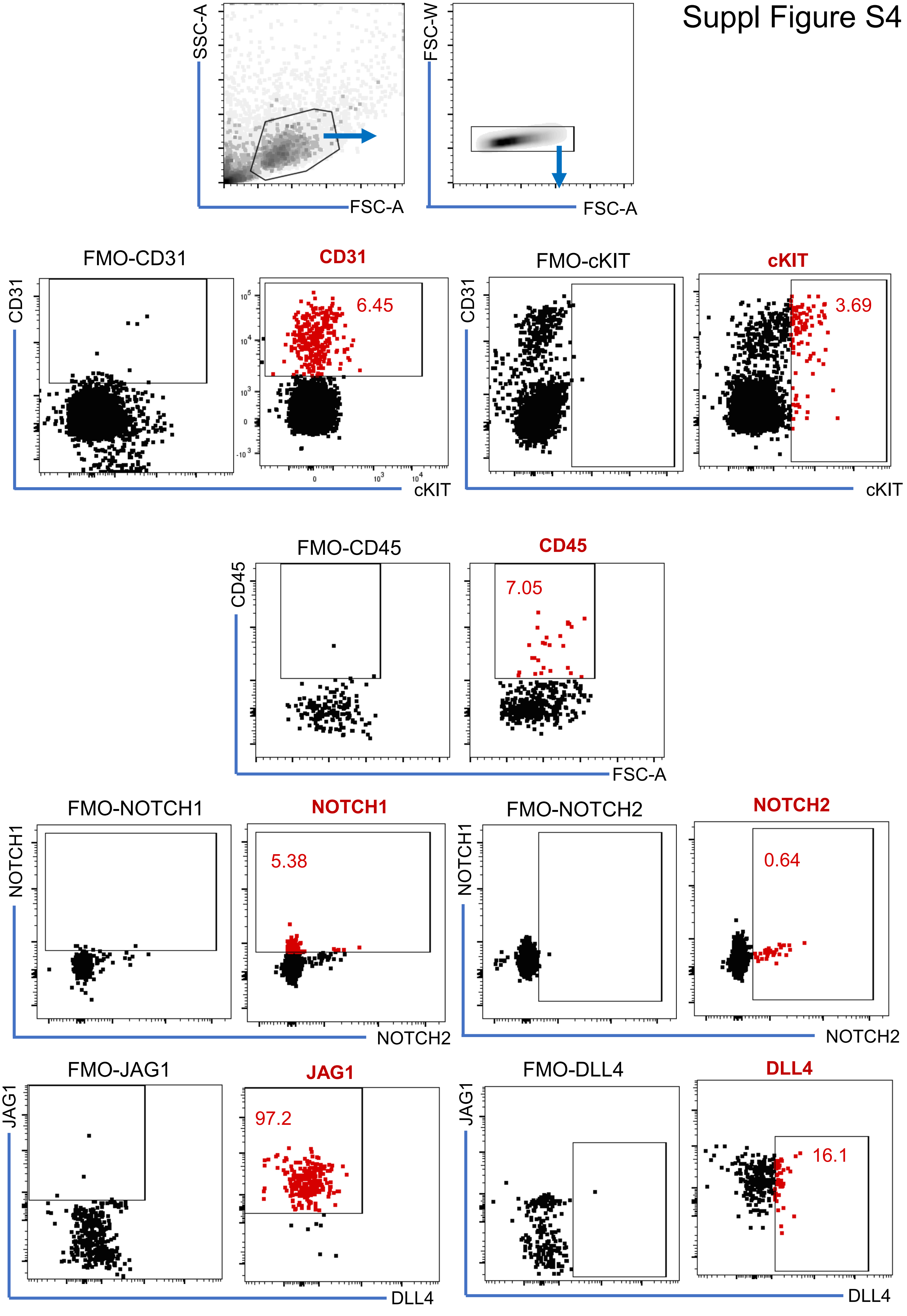

### Suppl Figure S5

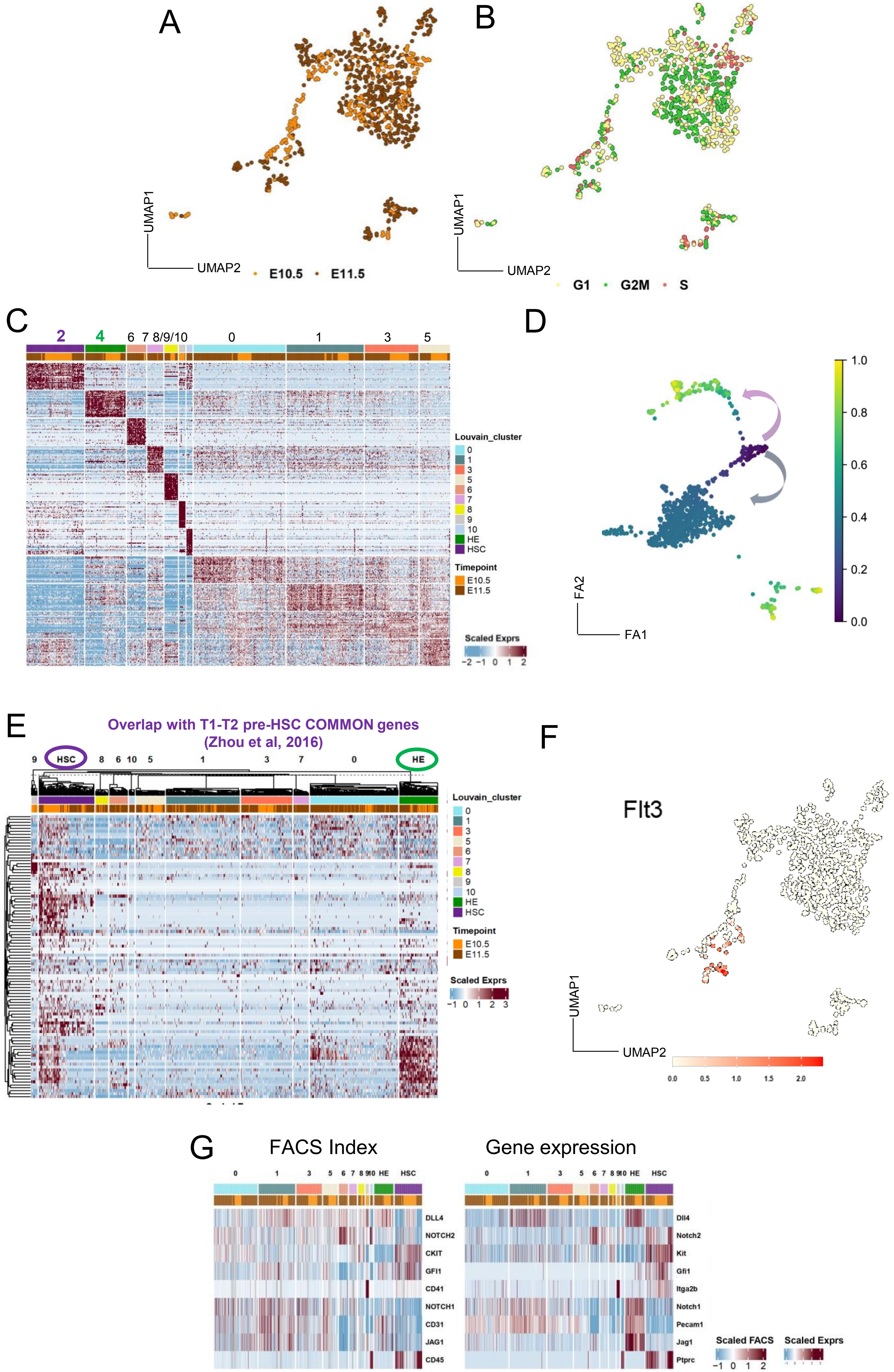

### Suppl Figure S6

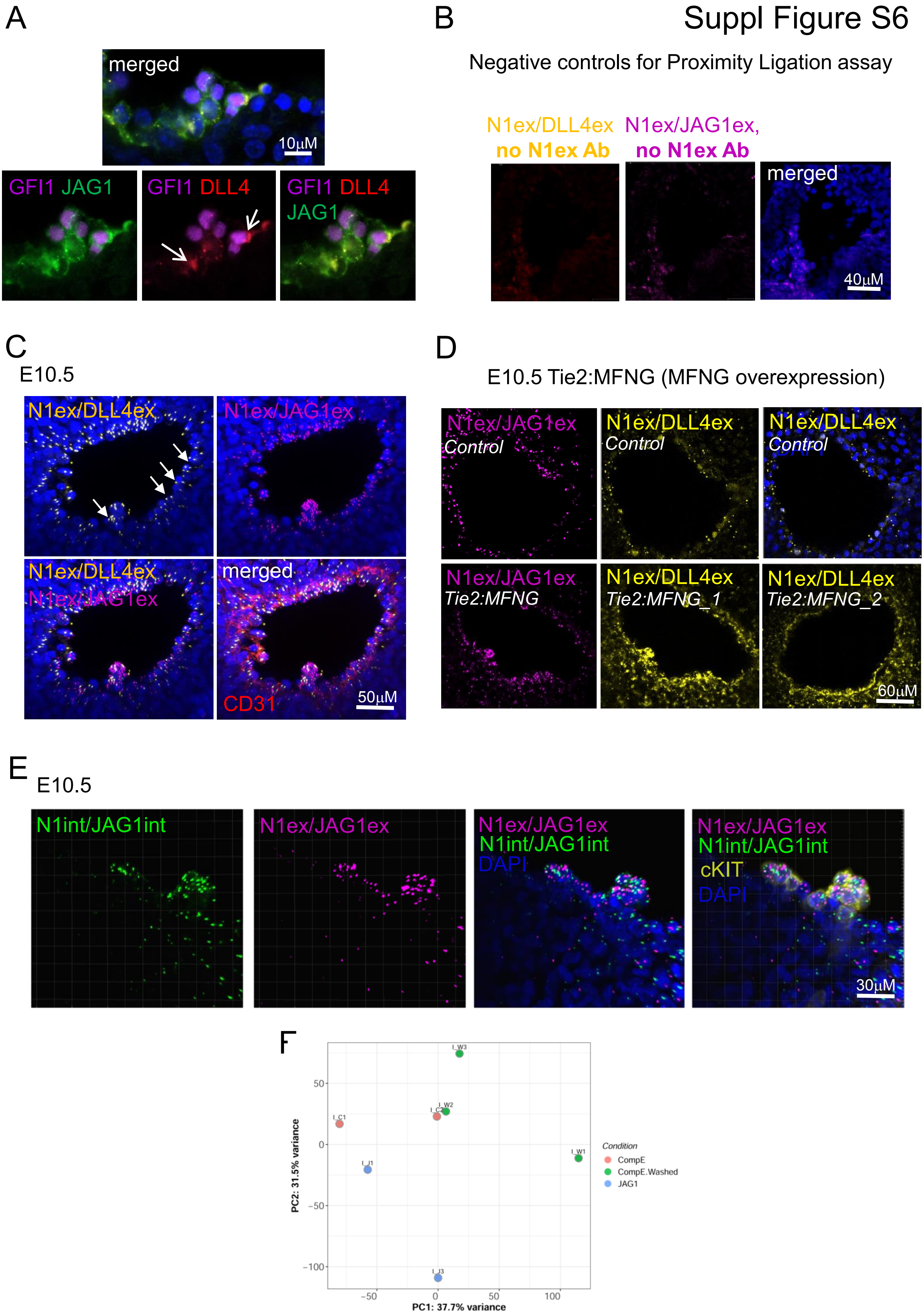

### Suppl Figure S7

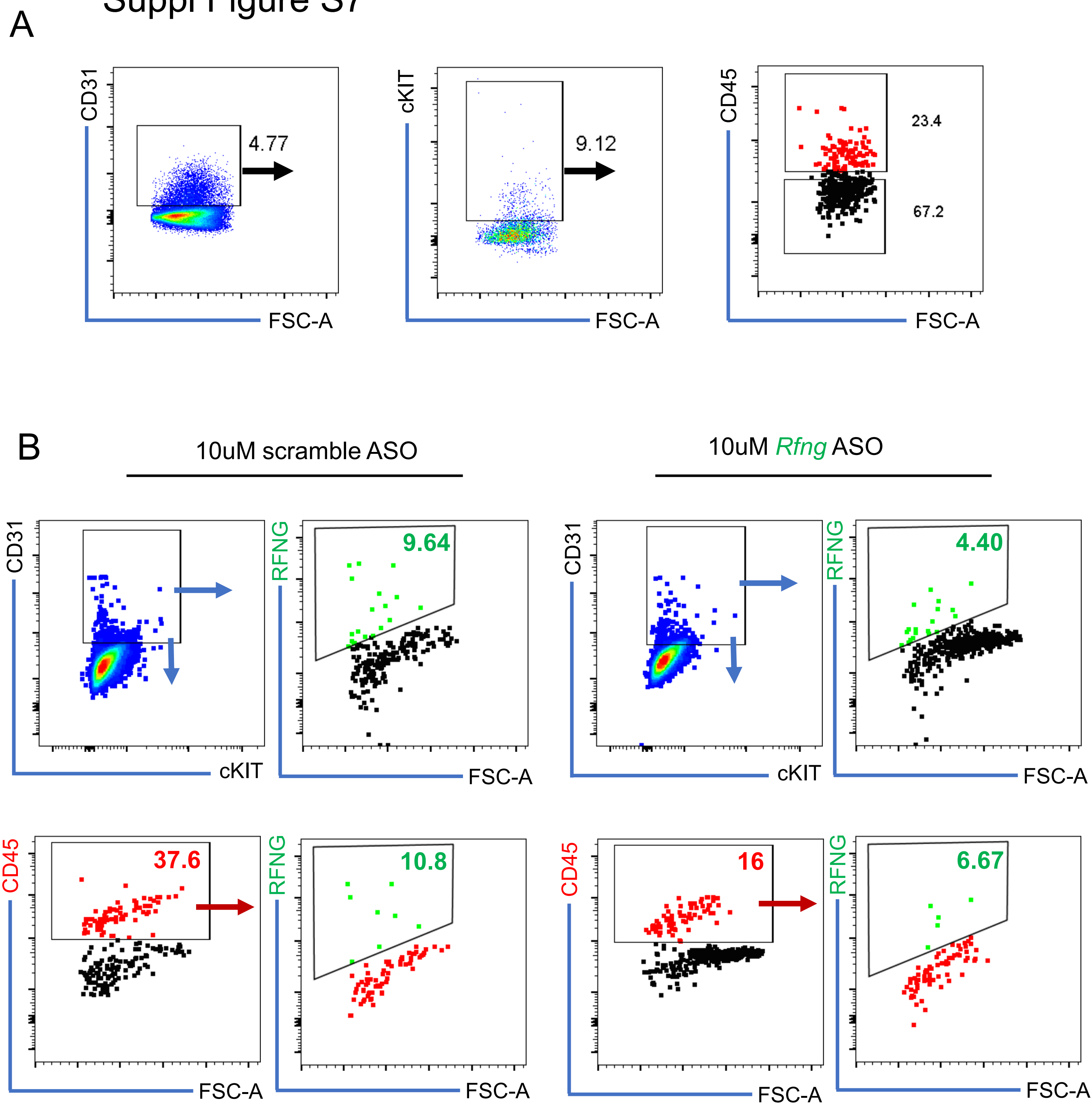
